# Predicting clinical drug response from model systems by non-linear subspace-based transfer learning

**DOI:** 10.1101/2020.06.29.177139

**Authors:** Soufiane Mourragui, Marco Loog, Daniel J. Vis, Kat Moore, Anna G. Manjon, Mark A. van de Wiel, Marcel J.T. Reinders, Lodewyk F.A. Wessels

## Abstract

Pre-clinical models have been the workhorse of cancer research for decades. While powerful, these models do not fully recapitulate the complexity of human tumors. Consequently, translating biomarkers of drug response from pre-clinical models to human tumors has been particularly challenging. To explicitly take these differences into account and enable an efficient exploitation of the vast pre-clinical drug response resources, we developed TRANSACT, a novel computational framework for clinical drug response prediction. First, TRANSACT employs non-linear manifold learning to capture biological processes active in pre-clinical models and human tumors. Then, TRANSACT builds predictors on cell line response only and transfers these to Patient-Derived Xenografts (PDXs) and human tumors. TRANSACT outperforms four competing approaches, including Deep Learning approaches, for a set of 15 drugs on PDXs, TCGA cohorts and 226 metastatic tumors from the Hartwig Medical Foundation data. For only four drugs Deep Learning outperforms TRANSACT. We further derived an algorithmic approach to interpret TRANSACT and used it to validate the approach by identifying known biomarkers to targeted therapies and we propose novel putative biomarkers of resistance to Paclitaxel and Gemcitabine.

## Introduction

The accumulation of somatic alterations on the genome and epigenome transforms healthy cells into malignant tumor cells. Although these alterations are required for tumor growth, they also confer vulnerabilities on tumor cells. Some well-known examples of such genetic vulnerabilities are the amplification of ERBB2 in breast cancer^1^, the BRAF^V600E^ mutation in skin melanoma^2^ or the BCR/ABL fusion in leukemia^3^. These vulnerabilities have been successfully exploited clinically by directing drugs against them. However, for the vast majority of cancer patients, no clear biomarkers exist. Hence, expanding our arsenal of accurate biomarkers would pave the way for personalized medicine, by identifying, for each patient, the most effective drug^4^.

In order to discover such biomarkers, pre-clinical models have been used extensively in the past decades, either in the form of cell lines, patient-derived xenografts (PDX) or organoids. This was partially fueled by the relative ease with which these model systems can be subjected to drug screening. This has led to break-through discoveries with broad clinical impact^5^. However, *Paul Valery*’s statement, “what is simple is always wrong; what is not, is unusable”^6^, also applies to these model systems. Specifically, their simplicity also confers weaknesses: the lack of a micro-environment in cell lines, and the absence of an immune system in cell lines, PDXs and organoids. These shortcomings are further amplified by culture artefacts^7,8^ that lead to a reduced clinical significance of these models^9,10^ and a high attrition rate in drug development^11^.

Computational approaches that correct for these differences are therefore needed to improve the identification of truly predictive biomarkers^12^. In the particular case of cancer, these approaches are divided into two distinct categories. In the first category, mechanistic models are developed on pre-clinical models and subsequently “humanized” to focus on the similarities between pre-clinical models and human tumors^13^. The second category approaches the problem in a statistical fashion. Using molecular profiles and drug screens from large-scale panels of pre-clinical models^14,15^, cell line drug response predictors are inferred^16–18^. The resulting models are then applied to predict the sensitivity of patients to certain drugs^19–21^. Although already promising, these approaches either do not take into account the fundamental differences between pre-clinical models and human tumors^22^, or only model these differences as a technical batch effect^19–21^. Recently, transfer learning and multi-task learning approaches have been developed to explicitly take these differences into account, either partially using tumor responses during training^23^, or solely based on pre-clinical labels^24^ while only modeling differences linearly.

We present **TRANSACT** (**T**umor **R**esponse **A**ssessment by **N**on-linear **S**ubspace **A**lignment of **C**ell-lines and **T**umors), a general framework for subspace-based transfer learning^25–29^ which enables the transfer of drug response predictors trained on a source domain (e.g. cell lines and PDXs) to a target domain (e.g. human tumors). TRANSACT employs the powerful mathematical framework of Kernel methods^30–35^ to capture both linear and non-linear molecular processes expressed in source and target domains. First, we demonstrate that, compared to existing methods^20,21,24^, modeling non-linearities using TRANSACT improves drug response prediction in PDXs after training on cell line responses only. We fix the hyperparameter controlling the degree of non-linearity on the PDX data and then employ TRANSACT to transfer predictors of drug response trained on cell lines to two human tumor datasets: primary tumors from TCGA and metastatic lesions from the Hartwig Medical Foundation (HMF). Specifically, we show a significant improvement in response prediction for five drugs in TCGA and three drugs in HMF, compared to state-of-the art competing approaches. Importantly, this performance improvement is attained without any training on data from the human tumors. We finally employ the interpretability of our approach to identify genes and pathways associated with drug response. We provide a full mathematical derivation of our algorithm, a complete reproducible pipeline and a fully open-source software package.

## Results

### TRANSACT: Generating non-linear manifold representations to transfer predictors of response from pre-clinical models to tumors

TRANSACT compares genomic signals contained in the source (e.g. pre-clinical models) and target (e.g. human tumors) datasets, and outputs a representation of processes that are present in both datasets. The nature of this representation depends on the similarity function *K* that characterizes the relationships between samples (Methods). Depending on the similarity function employed, various types of non-linear relationships can be modeled. For instance, in the case of a Gaussian similarity function, these non-linearities include constant, linear, second and higher-order interaction terms (Methods).

In a first step, TRANSACT computes the similarities between all samples (**Figure 1**A), yielding three matrices: *K*_*s*_, *K*_*t*_ and *K*_*st*_, containing the similarities between source samples, target samples, as well as source and target samples, respectively. Using an eigen-decomposition of *K*_*s*_ and *K*_*t*_, directions of maximal variance in the similarity space are computed using *Kernel PCA*^36^ (**Figure 1**B). This decomposition is performed independently on the source and the target spaces and yields two *importance score matrices*: one for the source non-linear principal components (NLPCs) and one for the target NLPCs. These importance scores are *not* the projected sample values –– they actually correspond to loadings, in the sense that they represent the geometric directions of the NLPCs in the sample similarity space (**Supp Figure 1**A). We then quantify the geometric differences between all pairs of NLPCs from the source and target sets by computing the cosine similarity matrix (**Figure 1**C). In a subsequent step (**Figure 1**D), we align these two sets of NLPCs by using the notion of *principal vector*s illustrated in **Supp Figure 1**B. These principal vectors (PV) are pairs of vectors – one from the source, one from the target – ranked by decreasing similarity: the top PVs correspond to geometrically similar factors, while bottom PVs are pairs of almost orthogonal vectors. We restrict further analysis to the most similar PVs based on a similarity of at least 0.5 (Supp. Material). We perform, for each selected pair of PVs, an interpolation between the source and the target vectors and select one intermediate representation that best balances the contribution of the source and the target signals (**Figure 1**E, **Supp Figure 1**C). These vectors, called *consensus features*, define the consensus space, into which we project the source (pre-clinical) and target (tumor) samples. This projection yields, for all source and target samples, consensus feature values that can be used as input to any machine learning model to build predictors of drug response (**Figure 1**G, Method). In the case of a linear similarity function, TRANSACT reduces to PRECISE^24^ (Subsection Supp. 8) and is fundamentally different from approaches such as Canonical Correlation Analysis (CCA)^37^ (Subsection Supp. 9).

**Figure 1.**
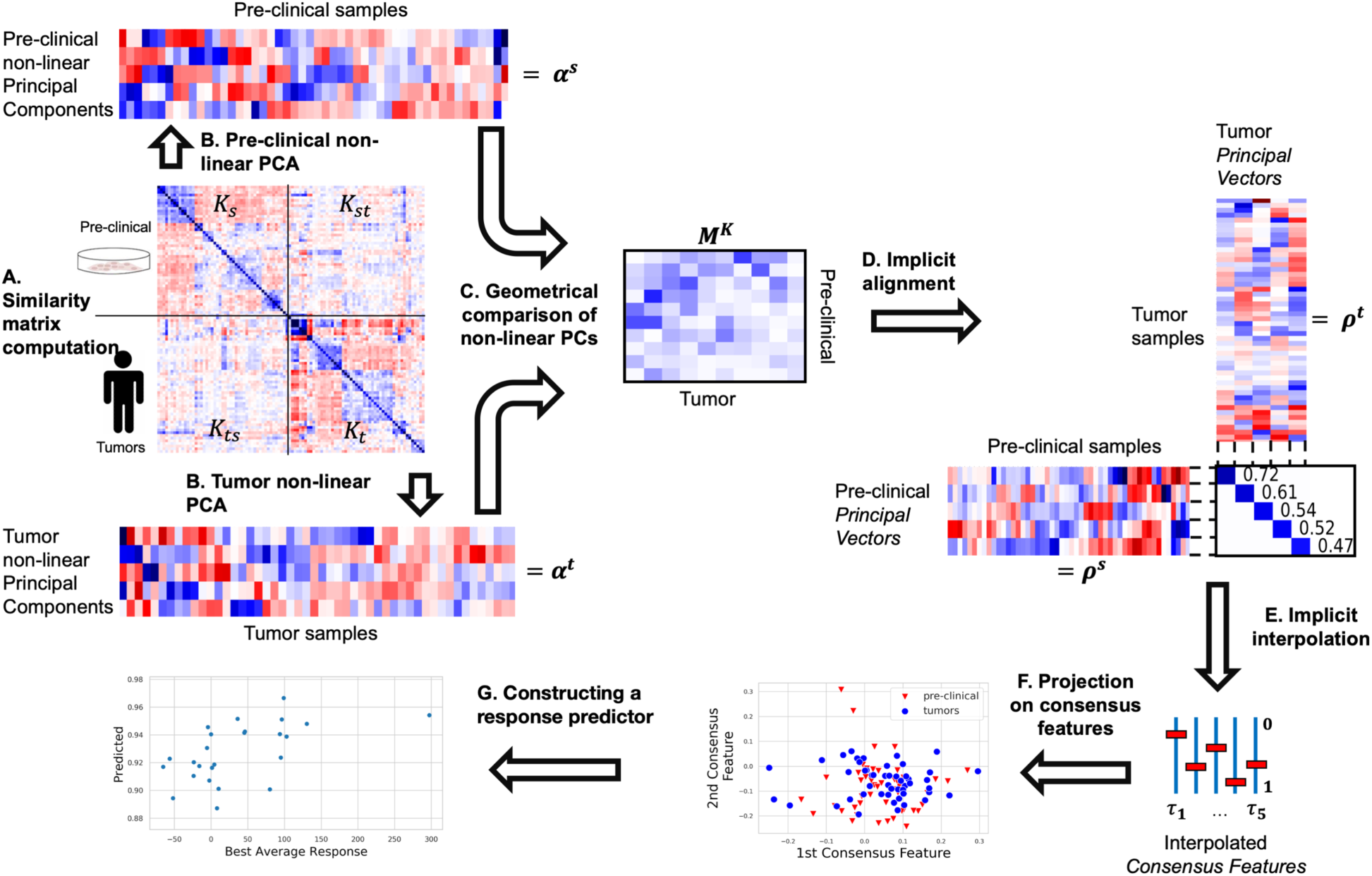
<u>TRANSACT: Generating non-linear manifold representations to transfer predictors of response from pre-clinical models to human tumors.</u> (**A**) Samples are compared using a **similarity function** yielding similarity matrices between pre-clinical models (source, *K*_*s*_), between tumors samples (target, *K*_*t*_) and between pre-clinical models and tumors (*K*_*st*_). (**B**) Using non-linear PCA, the pre-clinical and tumor similarity matrices are independently decomposed into **non-linear principal components** (NLPCs) geometrically represented by “sample importance scores” (**Supp Figure 1A**) that represent the importance of each sample in each NLPC (*α*^*s*^ and *α*^*t*^, for source and target space, respectively). (**C**) Geometrical comparison of pre-clinical and tumor NLPCs results in a **non-linear cosine similarity matrix *M***^*K*^. (**D**) Alignment of NLPCs using the notion of **principal vectors** (**Supp Figure 1B**). (**E**) Interpolation within each pair of vectors to select one vector per PV-pair that balances the effect of pre-clinical and tumor signals: the **consensus features** (**Supp Figure 1C**). (**F**) Projection of each tumor and pre-clinical sample on the consensus features to obtain **consensus scores**: scores that correspond to the activity of processes conserved between tumors and pre-clinical models. (**G**) Finally, these scores can be used as input to any predictive model, for instance to predict drug response based on these consensus scores.

### Non-linearities improve response prediction of predictors transferred from cell lines to patient-derived xenografts (PDXs)

When it comes to predicting drug response in one model system, it is known that inducing non-linearities can lead to improved performance^35^, although linear methods remain competitive^18,38^. We investigated whether the introduction of non-linearities in the computation of sample similarities resulted in improved response prediction of predictors trained on cell lines (source domain) and transferred to PDXs (target domain). Since gene expression is known to have predictive power comparable to other omics datasets combined^16,18,39,40^, we restricted our analysis to the expression of 1780 genes known to be related to cancer^41^. Using TRANSACT, we computed consensus features for cell lines (1049 cell lines from 26 different tissues) and all PDXs (399 samples from 5+ different tissues) (Methods). We projected the gene expression data of all cell lines and all PDXs onto these consensus features. We employed Elastic Net to train models of drug response with the projected cell line expression data as input and the drug response data as output, which is quantified as the AUC, i.e. the area under the drug response curve (Methods). We applied this trained predictor on the projected PDX expression data and compared the predicted response to the measured best average response by Spearman Correlation (**Figure 2**A). We made use of the standard Gaussian similarity function (Methods) to vary the level of non-linearity introduced. This similarity function is characterized by a single scaling factor *γ*, whose size is directly proportional to the proportion of non-linearity introduced (**Figure 2**B). We studied the predictive performance in PDXs for seven different values of *γ*, ranging from a set of consensus features with an almost purely linear (*γ* = 1 × 10^−5^) to an almost purely non-linear composition (*γ* = 1 × 10^−2^). We compared the performance of TRANSACT to two approaches that do not perform domain adaptation: *Elastic Net*^22^ and *Deep Learning* regression (DL). We further compared it to two state-of-the-art domain adaptation approaches: ComBat batch correction followed by Deep Learning regression (*ComBat + DL*)^21^ and *PRECISE*^24^, a linear domain adaptation approach. All models were trained to predict response to seven different drugs (Erlotinib, Cetuximab, Gemcitabine, Afatinib, Paclitaxel, Trametinib and Ruxolitinib) for which we had response data available for both PDX models and cell lines (**Figure 2**C-I). For these seven drugs we observe, in general, a clear improvement of approaches that employ domain-adaptation over those that do not, indicating a necessity to correct the input signal when moving from the source to the target domain. When comparing *Elastic Net* to *Deep Learning*, we observe that modeling non-linearities results in a better transfer for five drugs. We further observe an improvement of TRANSACT over *ComBat+DL*, suggesting that the correction required from cell lines to PDXs is more complicated than correcting for a technical batch effect. We observe that the introduction of non-linearity with TRANSACT generally increases the predictive performance on PDXs when compared to a linear similarity function (*PRECISE*). Specifically, we observe for several drugs that the predictive performance increases with the scaling factor until a maximal performance is reached (*γ* = 10^−4^ for Erlotinib, Cetuximab and Afatinib and *γ* = 10^−3^ for Gemcitabine, Paclitaxel and Trametinib), after which the predictive performance drops. We therefore decided to fix the scaling factor to the average of these two values (*γ* ^*^ = 5 × 10^−4^) and employ the associated consensus space to transfer the predictors of response to the tumor samples. As a further check, we analyzed the properties of the consensus space obtained using *γ* ^*^. We observe a concentration of the offset contribution in the top consensus features and an increasing proportion of non-linear term contribution to lower order features (**Supp Figure 6**C). The UMAP^42^ projection of the consensus features shows a clear co-clustering of cell lines and PDXs of the same tissue (**Supp Figure 6**D).

**Figure 2.**
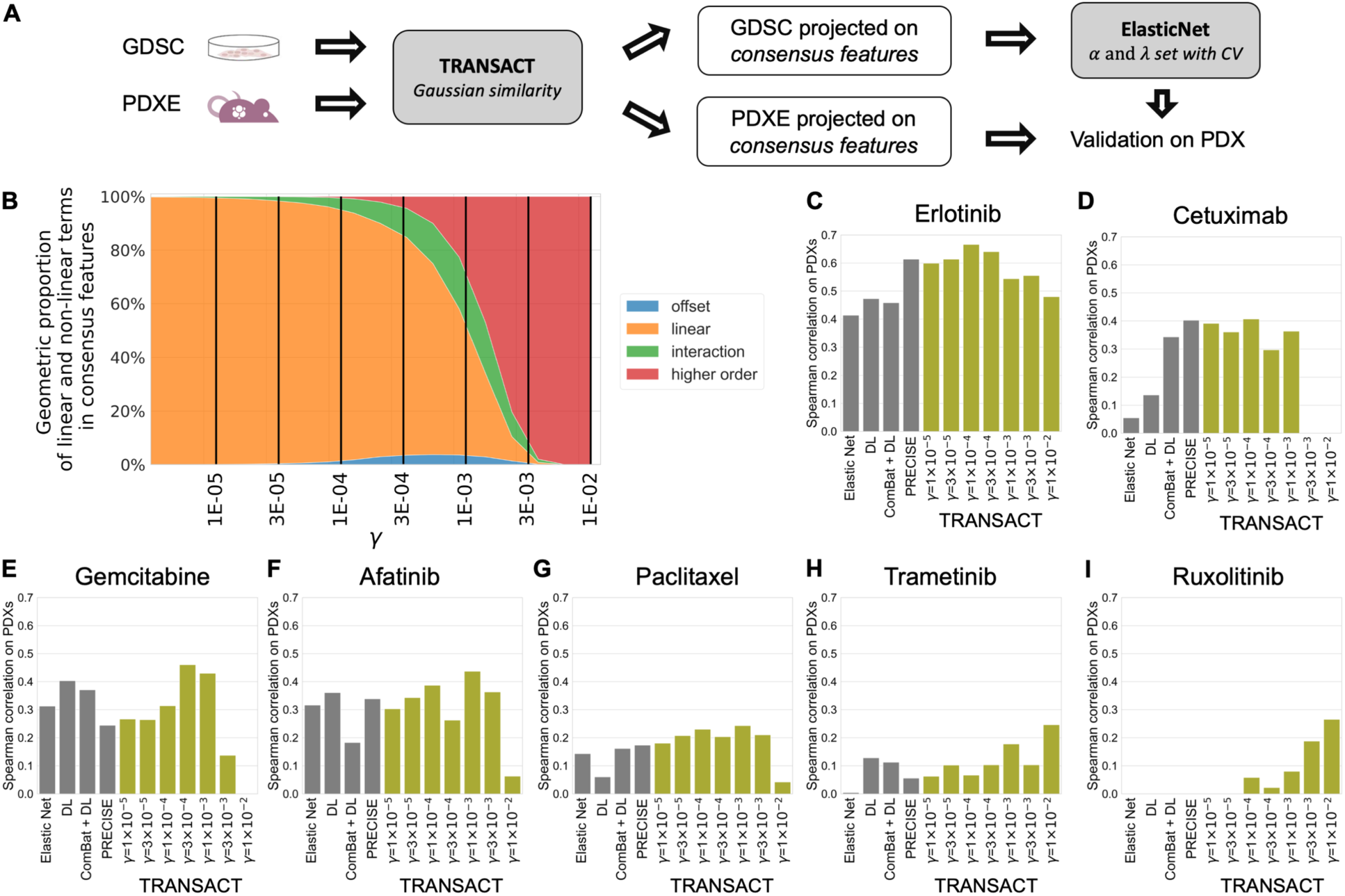
<u>Impact of modeling non-linearities for drug response prediction transfer from cell lines to PDXs.</u> (**A**) Main workflow of the prediction on PDXs. Using cell lines and PDX gene expression, we compute consensus features and project each dataset onto these. We then trained a regression model (Elastic Net) on cell lines projected scores to predict response, measured as Area Under the drug response Curve (AUC). Finally, we used this regression model to predict drug response in PDXs and correlate the predicted AUC to the known best average response. (**B**) Proportion of non-linearities induced by the Gaussian similarity function as a function of the scaling factor *γ*. For different values of *γ*, we computed the average contribution over all consensus features of offset, linear, interaction and higher order features (Methods). Offset is here to be understood as the exponential of the squared depth and does not correspond to a constant term. We finally evaluated the response prediction on the PDX models for different values of *γ*, and for four competing approaches: two methods (*ElasticNet* and *Deep Learning*) that do not model the transfer, and two transfer learning methods (ComBat followed by Deep Learning, and *PRECISE*). We report results for Erlotinib (**C**), Cetuximab (**D**), Gemcitabine (**E**), Afatinib (**F**), Paclitaxel (**G**), Trametinib (**H**) and Ruxolitinib (**I**). Concordance on PDX is measured as the Spearman correlation between predicted AUC and Best Average Response.

### Consensus features between cell lines (GDSC) and human tumors conserve primary tumor information

With the scaling factor (*γ*) calibrated on PDX models, we moved to the clinical setting to investigate domain adaptation between cell lines and two different human tumor datasets: primary tumors from TCGA and metastatic lesions from the HMF. We selected 30 consensus features in the GDSC-TCGA analysis (**Supp Figure 8**) and 20 in the GDSC-HMF analysis (**Supp Figure 9**). We arrived at these numbers by first selecting NLPCs based on the inflexion point of the cumulative eigenvalues, and subsequently only retaining PVs with a similarity above 0.5. We observe that the consensus features computed between GDSC and TCGA (**Figure 3**A) and between GDSC and HMF (**Figure 3**B) show the same proportion of non-linearity with a concentration of offset and linearities in the top consensus features.

**Figure 3.**
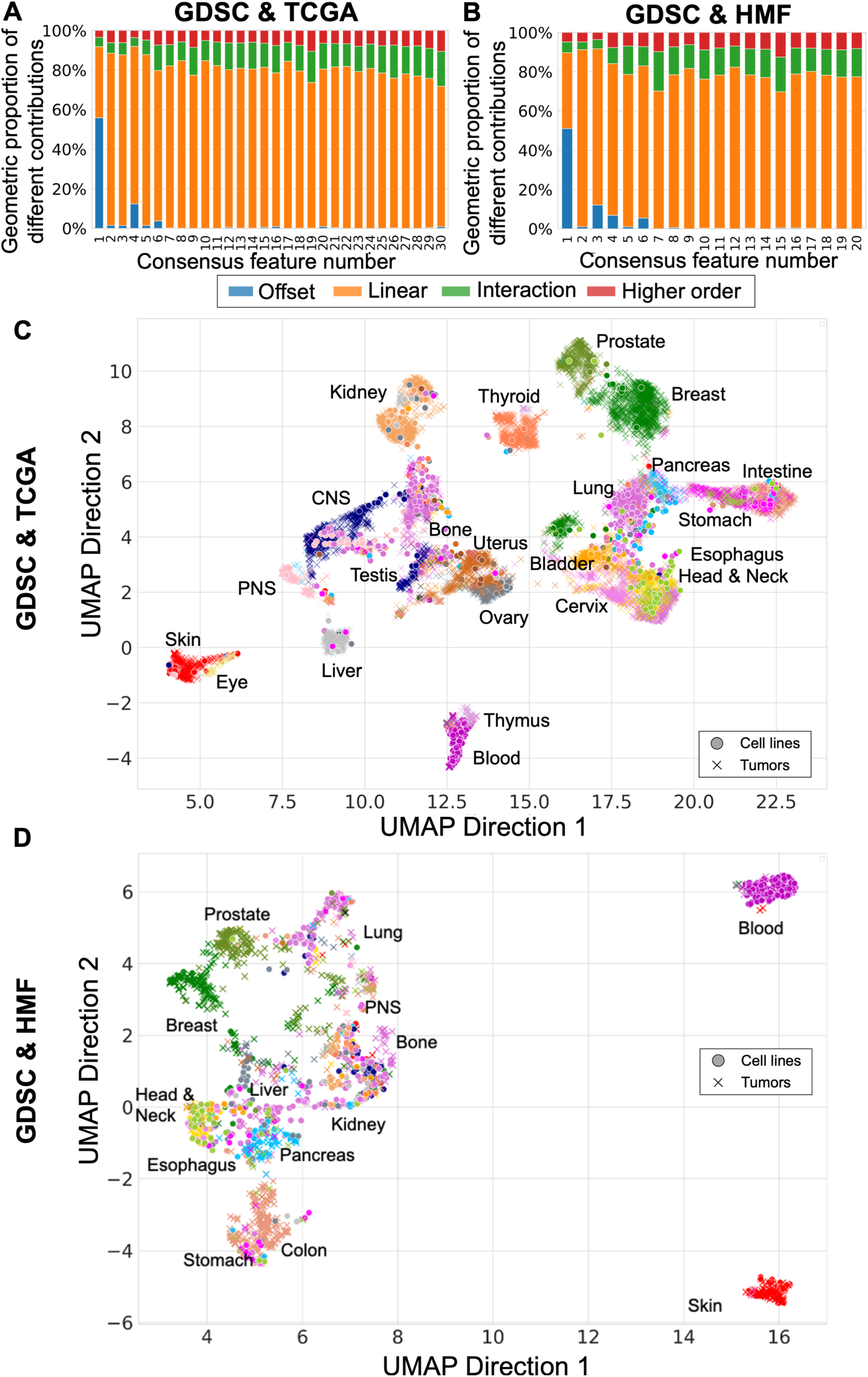
<u>Pan-cancer consensus features between cell lines and tumors conserve tissue type information</u>. We used cell lines (GDSC) as source data and computed two sets of consensus features with two different target datasets: primary tumors (TCGA, A and C) and metastatic lesions (HMF, B and D). (**A**) Proportion of linear and non-linear contributions to each of the 30 GDSC-to-TCGA consensus features. (**B**) Proportion of linear and non-linear contributions to each of the 20 GDSC-to-HMF consensus features. (**C**) UMAP plot of primary tumors (TCGA, 21 tissues) and cell lines (GDSC, 22 tissues) projected on the consensus features, using the same parameters as selected in **Figure *2***. (**D**) UMAP plot of metastatic lesions (HMF) and cell lines, colored by primary tissue for both HMF and GDSC. For both UMAP plots, the full legend can be found in **Supp Figure 10**A.

In order to visualize the structure retained in the consensus space, we embedded our consensus scores into a 2D space using UMAP^42^. We observed that primary tumors cluster together based on their tissue type (**Figure 3**C). Cell lines, however, show different behaviors – most do cluster with the tumors from a similar tissue of origin, while a group of cell lines cluster together and away from the tumors, regardless of their tissue of origin, as observed in previous studies^43^. To quantify the degree of co-clustering of cell lines and tumors, we compared distances between tumors and cell lines from similar and non-similar tissues, and observed, as expected, a higher similarity between tumors and cell lines from the same tissue (**Supp Figure 10**C). Metastatic lesions show a weaker clustering based on the primary tumor’s tissue of origin (**Supp Figure 10**D-E). This is not unexpected, as the expression profiles are derived from biopsy sites distant from the primary tissue. Of particular interest, we observe the existence of a hematopoietic cell-line cluster that co-clusters with metastatic samples from various biopsy sites. Most of these tumor samples (7 out of 12 samples) are lymph node metastases and most likely display a hematopoietic expression profile due to blood infiltration in the samples (**Supp Figure 10**B).

### Consensus features increase transfer of response predictors from cell lines to primary tumors and metastatic lesions

To further validate our approach, we transferred response predictors from cell lines to the TCGA and HMF collections of human tumors. First, we projected the GDSC and TCGA expression data onto the GDSC-TCGA consensus features. Then we trained, for each drug, a regression model using solely the cell line response data (measured as AUC). These drug-specific regression models were then used to predict response on the projected TCGA data. Finally, we compared the predicted response to the known categorical clinical responses using a one-sided Mann-Whitney test and computed the corresponding effect size. We trained models for seventeen different drugs (**Table 1**A). We compared the performance of TRANSACT to the performance obtained by four competing approaches (*Elastic Net, DL, ComBat+DL* and *PRECISE*) (**Table 1**A, Methods). For the Deep Learning approaches (*DL* and *ComBat+DL*), we selected the architecture and hyper-parameters for each drug by 5-fold cross-validation on GDSC (Methods). We subsequently trained 50 models with different and independent initializations (Method) and reported the median performance obtained on TCGA. *Elastic Net* and *PRECISE* obtain significant associations for three and six drugs, respectively, but neither approach ever outperforms all other approaches. *ComBat+DL* and *DL* achieve significant associations for five and eight drugs, respectively – however, both approaches outperform all others for only three and one drugs, respectively. By contrast, TRANSACT achieves significant associations for seven drugs and outperforms all other approaches for five drugs. For both deep learning approaches, we observe an important dependency on the network initialization (**Supp Figure 11**A, **Supp Figure 12**A). More importantly, we observe no correlation between the training error achieved on the source domain and the prediction accuracy on the target domain, making it impossible to select a proper initialization solely based on the source domain performance (**Supp Figure 11**B, **Supp Figure 12**B). Results obtained with TRANSACT, on the contrary, do not depend on a random initialization.

**Table 1.** <u>Results of TRANSACT compared to four competing approaches –</u> *<u>Elastic Net, Deep Learning, ComBat+DL</u>*<u>, and</u> *<u>PRECISE</u>* <u>– for 17 drugs on TCGA and 5 drugs on HMF.</u> For each drug, we train 5 predictors and compare in each scenario the predicted AUC to the known clinical response using one-side Mann-Whitney test. The clinical response corresponds to the following division: Responders and Non-Responders for TCGA, PR and PD for HMF (Methods). For each predictor, we report the p-value and the effect size (Area under the ROC, effect size associated with the Mann-Whitney test) under bracket. Bold cells correspond to significant association (pval < 0.05). Red cells correspond to significant associations with the largest effect size across the three methods.

| A. GDSC-to-TCGA |  |  |  |  |  |  |  |  |
| --- | --- | --- | --- | --- | --- | --- | --- | --- |
| GDSC |  | TCGA |  | p-val [effect-size] on TCGA |  |  |  |  |
| Drug | Samples | Drug | Samples | Elastic Net | Deep Learning | DL + ComBat | PRECISE | TRANS ACT |
| Afatinib | 800 | Trastuzumab | 16 | 0.089<br>[0.82] | 0.034<br>[0.93] | 0.048<br>[0.89] | 0.026<br>[0.96] | 0.016<br>[1.00] |
| Bleomycin | 856 | Bleomycin | 53 | 0.15<br>[0.63] | 0.19<br>[0.61] | 0.41<br>[0.53] | 0.11<br>[0.66] | 0.091<br>[0.67] |
| Cetuximab | 868 | Cetuximab | 19 | 0.12<br>[0.67] | 0.52<br>[0.50] | 0.39<br>[0.54] | 0.45<br>[0.52] | 0.077<br>[0.70] |
| Cisplatin | 764 | Cisplatin | 308 | 1.1E-6<br>[0.69] | 1.8E-4<br>[0.64] | 1.1E-2<br>[0.59] | 2.2E-5<br>[0.66] | 3.9E-7<br>[0.70] |
| Cisplatin | 764 | Carboplatin | 166 | 0.042<br>[0.58] | 0.10<br>[0.56] | 0.53<br>[0.50] | 0.024<br>[0.59] | 0.0035<br>[0.63] |
| Cyclophosphamide | 747 | Cyclophosphamide | 102 | 0.57<br>[0.48] | 0.67<br>[0.45] | 0.89<br>[0.35] | 0.11<br>[0.65] | 0.64<br>[0.46] |
| Docetaxel | 665 | Docetaxel | 102 | 0.62<br>[0.48] | 0.27<br>[0.54] | 0.54<br>[0.49] | 0.78<br>[0.46] | 0.11<br>[0.58] |
| Doxorubicin | 871 | Doxorubicin | 101 | 0.81<br>[0.45] | 0.0055<br>[0.66] | 0.60<br>[0.48] | 0.99<br>[0.32] | 0.69<br>[0.47] |
| Etoposide | 880 | Etoposide | 84 | 0.21<br>[0.58] | 0.0032<br>[0.76] | 0.0076<br>[0.73] | 0.0071<br>[0.73] | 0.027<br>[0.68] |
| 5-Fluorouracil | 801 | Fluorouracil | 186 | 0.22<br>[0.54] | 0.72<br>[0.47] | 0.79<br>[0.46] | 0.80<br>[0.46] | 0.35<br>[0.52] |
| Gemcitabine | 752 | Gemcitabine | 156 | 0.25<br>[0.53] | 0.021<br>[0.59] | 0.0057<br>[0.62] | 0.029<br>[0.59] | 0.0047<br>[0.62] |
| Irinotecan | 796 | Irinotecan | 25 | 0.81<br>[0.39] | 0.42<br>[0.53] | 0.68<br>[0.44] | 0.54<br>[0.49] | 0.49<br>[0.51] |
| Oxaliplatin | 724 | Oxaliplatin | 66 | 0.38<br>[0.52] | 0.022<br>[0.65] | 0.0099<br>[0.68] | 0.028<br>[0.64] | 0.035<br>[0.64] |
| Paclitaxel | 753 | Paclitaxel | 160 | 0.032<br>[0.59] | 0.025<br>[0.60] | 0.28<br>[0.53] | 0.10<br>[0.56] | 0.0042<br>[0.63] |
| Pemetrexed | 898 | Pemetrexed | 38 | 0.16<br>[0.60] | 0.39<br>[0.53] | 0.55<br>[0.49] | 0.16<br>[0.59] | 0.24<br>[0.57] |
| Temozolomide | 746 | Temozolomide | 96 | 0.56<br>[0.49] | 0.29<br>[0.55] | 0.47<br>[0.51] | 0.59<br>[0.48] | 0.19<br>[0.58] |
| Vinorelbine | 746 | Vinorelbine | 30 | 0.31<br>[0.57] | 0.043<br>[0.72] | 0.053<br>[0.71] | 0.19<br>[0.61] | 0.42<br>[0.53] |
| B. GDSC-to-HMF |  |  |  |  |  |  |  |  |
| GDSC |  | HMF |  | p-val [effect-size] on HMF |  |  |  |  |
| Drug | Samples | Drug | Samples | Elastic Net | Deep Learning | DL + ComBat | PRECISE | TRANS ACT |
| Afatinib | 800 | Trastuzumab | 25 | 0.18<br>[0.70] | 0.069<br>[0.81] | 0.23<br>[0.67] | 0.021<br>[0.93] | 0.032<br>[0.89] |
| Irinotecan | 796 | Irinotecan | 67 | 0.060<br>[0.73] | 0.060<br>[0.73] | 0.060<br>[0.73] | 0.10<br>[0.69] | 0.020<br>[0.80] |
| Cisplatin | 764 | Carboplatin | 64 | 0.23<br>[0.58] | 0.051<br>[0.67] | 0.094<br>[0.64] | 0.59<br>[0.67] | 0.0045<br>[0.78] |
| 5-Fluorouracil | 801 | Fluorouracil | 65 | 0.065<br>[0.68] | 0.12<br>[0.64] | 0.16<br>[0.62] | 0.21<br>[0.59] | 0.24<br>[0.58] |
| Paclitaxel | 753 | Paclitaxel | 45 | 0.56<br>[0.48] | 0.32<br>[0.56] | 0.39<br>[0.54] | 0.39<br>[0.53] | 0.061<br>[0.70] |
| Gemcitabine | 752 | Gemcitabine | 50 | 0.039<br>[0.71] | 0.014<br>[0.76] | 0.039<br>[0.71] | 0.0089<br>[0.78] | 0.0042<br>[0.81] |

For the HMF data, we repeated the steps above, while employing the GDSC-HMF consensus features as well as the HMF and GDSC expression and response data. We trained models for Irinotecan, Carboplatin (using Cisplatin GDSC response), Trastuzumab (using Afatinib GDSC response), Gemcitabine, Paclitaxel and 5-Fluorouracil. Across all approaches, we observe a significant association between the predicted AUC and clinical responses for four of the six drugs (Irinotecan, Carboplatin, Trastuzumab and Gemcitabine) (**Table 1**B, **Figure 4**A). *PRECISE* reaches significance for two drugs, whereas *Elastic Net, DL* and *Combat+DL* reach significance for a single drug. *TRANSACT*, by contrast, outperforms *PRECISE* on three of the five drugs, and *ComBat + DL, Deep Learning* and *Elastic Net* on four drugs. *TRANSACT* does not obtain a significant association for Paclitaxel. We do, however, observe a trend that indicates a positive association between predicted and clinical response. In contrast, all competing approaches fail to achieve any association. **Figure 4**A provides a summary of the performance of TRANSACT and two state-of-the-art approaches, *PRECISE* and *Combat + DL*, on both TCGA and HMF cohorts. Finally, for the HMF cohort, we also observe a large variability in predictions of the deep learning approaches as a result of network initialization (**Supp Figure 11**C, **Supp Figure 12**C). As on the TCGA we do not observe any relationship between the training error on the source domain data and the predictions on the target domain data (**Supp Figure 11**D, **Supp Figure 12**D).

**Figure 4.**
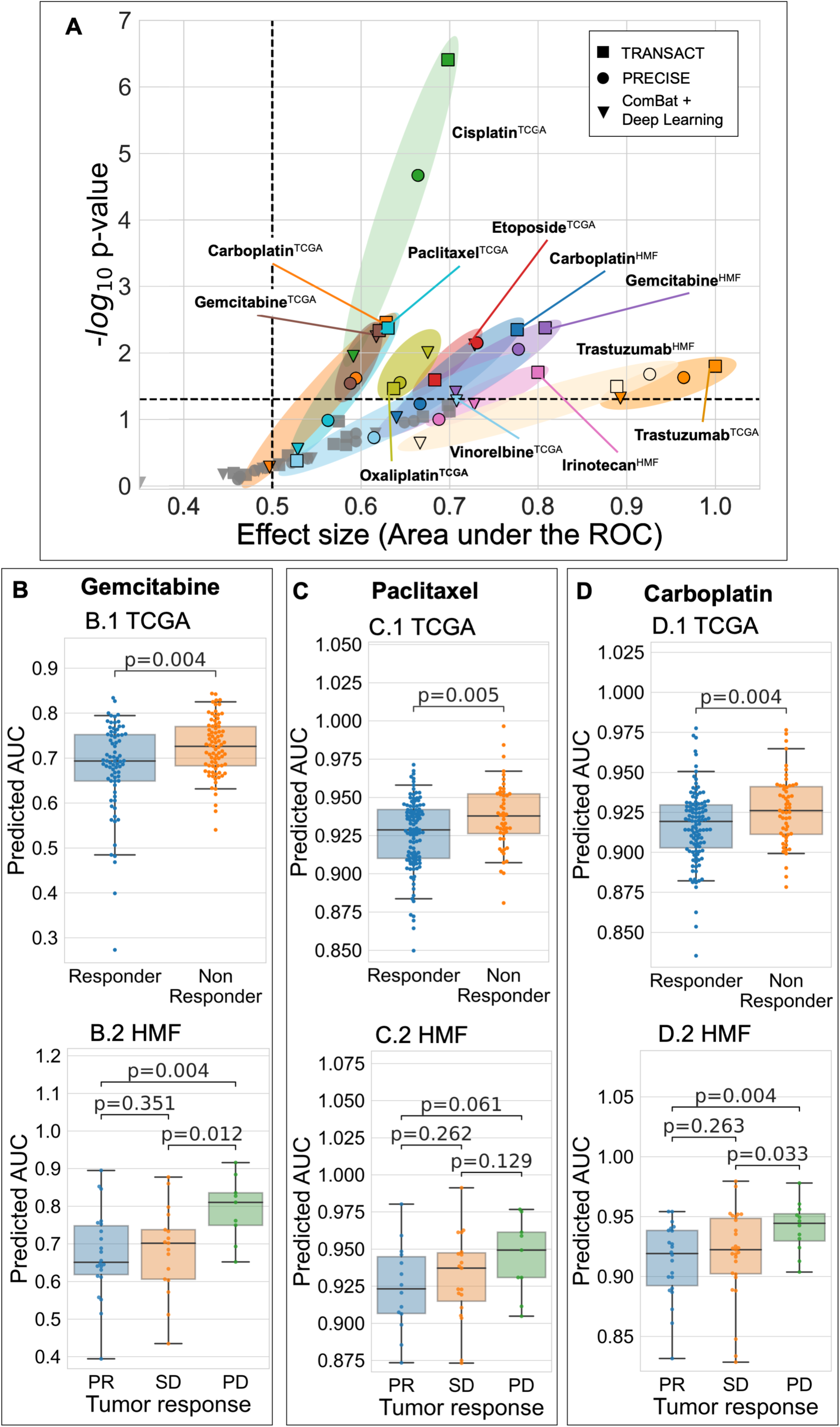
<u>Consensus features improve response prediction in patients.</u> (**A**) We used consensus features computed between GDSC and each tumor dataset to train a predictor for 17 drugs on TCGA and 5 drugs on HMG, using GDSC response only. We then predicted the Area Under the drug response Curve (AUC) for each drug and compared this predicted value to the observed clinical response using a one-sided Mann-Whitney test (**Table 1**). We performed the same prediction tasks using two state-of-the-art approaches (*ComBat + Deep Learning* and *PRECISE*, **Method**) and summarized here the associations for the 3 × 22 predictors in a Volcano plot, with the corresponding p-values on the y-axis and the effect-sizes on the x-axis. (**B**) Results for the two Gemcitabine predictors. (**B**.1) For the Gemcitabine predictor on TCGA, we compared predicted AUC for each patient to the known clinical response. (**B**.2) Results for Gemcitabine predictor on HMF. (**C**.1) Results for Paclitaxel predictor on TCGA. (**C**.2.) Results for Paclitaxel predictor on HMF. (**D**.1) Results for Carboplatin predictor on TCGA. (**D**.2.) Results for Carboplatin predictor on HMF.

Two drugs, already standing out from the GDSC-PDX analysis, are of particular interest. We first observe that Gemcitabine is consistently better predicted by *TRANSACT* than by any other approaches (**Figure 4**B, **Table 1**A). When it comes to Paclitaxel (**Figure 4**C, **Table 1**A), *TRANSACT* shows a clear improvement over competing approaches in TCGA. *TRANSACT* is also the only method to recover a positive association on HMF, although the association is not significant. Finally, for Carboplatin (**Figure 4**C, **Table 1**A), we observe that *TRANSACT* outperforms all approaches on TCGA, and on HMF. When taking the TCGA and HMF results together – and noting that there is overlap in the drugs employed in these two cohorts – it is interesting to observe that the best competing approach is *Deep Learning. Deep Learning* achieves significant associations in 9 of the 23 comparisons while *ComBat+DL* reaches significance in 6 trials. Based on this metric, *TRANSACT* performs best, reaching significance in 11 of 23 trials. Furthermore, *Deep Learning* outperforms all other approaches in only 3 comparisons, while *TRANSACT* delivers increased performance in 8 comparisons. When also taking the performance on the PDX models into account, we can safely conclude that *TRANSACT* is more accurate in transferring drug response predictors.

### Interpretability of consensus features confirms known mechanisms for targeted therapies and unveils potential biomarkers of sensitivity for cytotoxic drugs

We finally made use of the interpretability of our approach to mechanistically validate our predictors (Methods). We first validated targeted therapies with documented modes of action. We started with the TRANSACT predictor of response for Afatinib, a small molecule inhibitor of the EGFR family, which includes HER2 (**Figure 5**A). We performed a gene set enrichment analysis of the linear terms that constitutes to 80% of the predictor (**Supp. Table 1**). Most enriched gene sets are related to breast cancer subtypes as defined by *Charafe and colleagues*^44^ where, contrary to the definition based on the intrinsic breast cancer subtypes, the Luminal subtype contains both ER+ and HER2+ tumors. The top ranked gene set amongst the genes associated with sensitivity (genes with a negative coefficient in the predictor) are genes associated with the “Luminal” subtypes (FDR < 0.001). Conversely, genes associated with resistance (genes with a positive coefficient in the predictor) show enrichment for the “Mesenchymal” molecular signatures, shared by basal and mesenchymal subtypes. This corresponds with HER2 negative samples, which is in line with our expectation as absence of the drug target would indicate lack of response. Similarly, in the TRANSACT response predictor for Gefitinib (EGFR inhibitor) the genes constituting the linear portion and associated with sensitivity (negative predictor coefficients) show an enrichment for genes downregulated in Gefitinib resistant tumors (**Figure 5**B, **Supp. Table 2**).

**Figure 5.**
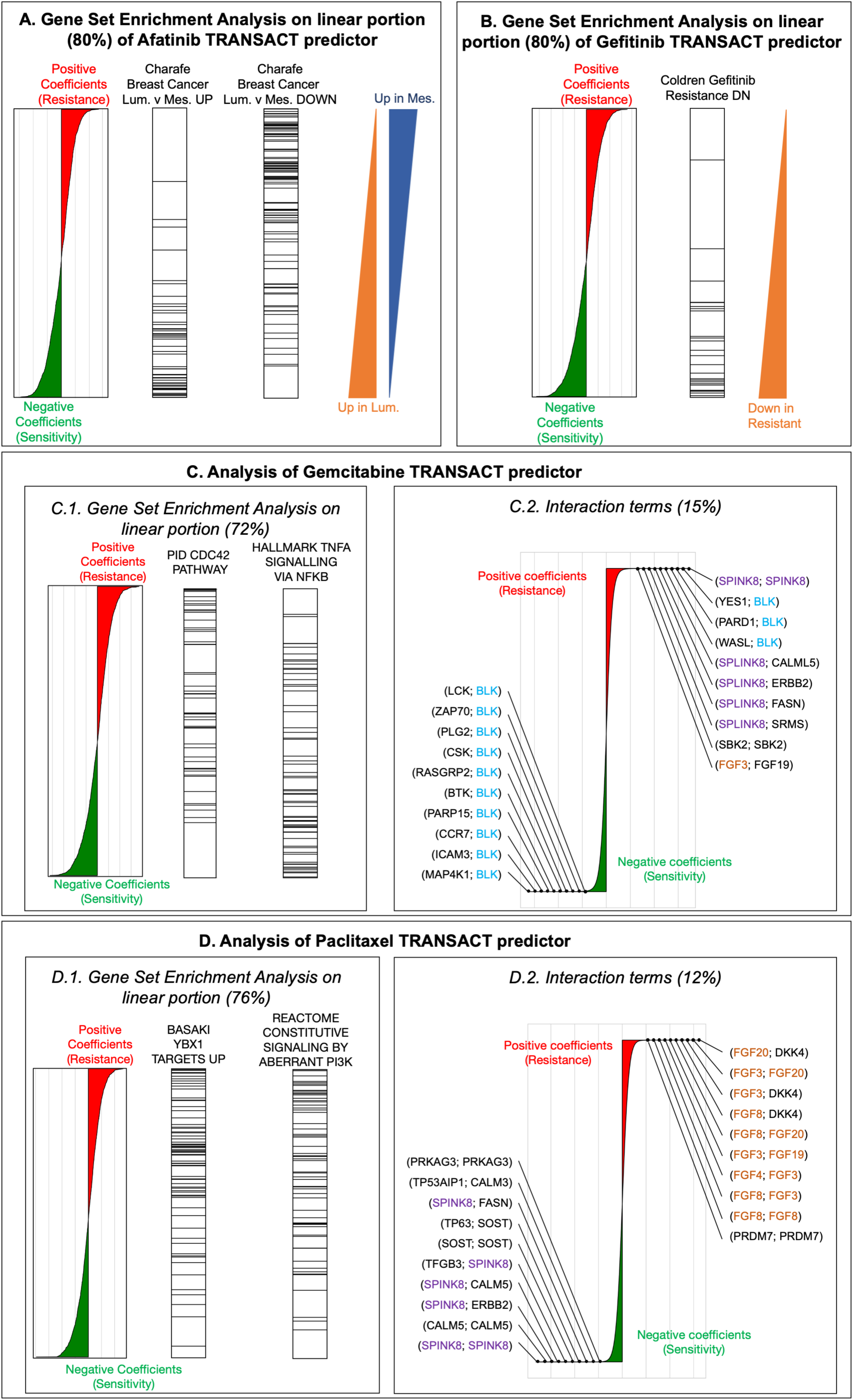
<u>Interpretability of TRANSACT consensus features recapitulates modes of action for Afatinib and Gefitinib, and highlights mechanisms of sensitivity and resistance to Gemcitabine and Paclitaxel.</u> (**A**) For the Afatinib (HER-2 protein kinase inhibitor) response model trained on GDSC samples projected on GDSC-to-TCGA consensus features, we show the contribution of each gene to the predictor (left): positive (negative) weights in the predictor indicate that high (low) expression of the genes leads to resistance (sensitivity) represented by larger (smaller) AUCs. We performed a PreRanked gene set enrichment analysis (GSEA) on these weights and highlight genes associated to two significantly enriched gene sets: “Charafe [*…*] Luminal vs Mesenchymal UP” (center-left) and “Charafe [*…*] Luminal vs Mesenchymal DOWN” (center-right). The two triangles on the right illustrate these two gene-set distributions. (**B**) Interpretation of the linear part of the Gefitinib (EGFR inhibitor) model. We display gene weights ordered by contribution (left) and the ranks of genes known to be down regulated in Gefitinib resistant tumors (right). (**C**) Interpretation of Gemcitabine predictor. (*C*.*1*.) We display the positions of genes associated to two significantly enriched gene sets: “CDC42 pathway”, associated to positive weights (resistance), and “TNF*α*-signalling”, associated to negative weights (sensitivity). (*C*.*2*.) Interaction terms make 15% of Gemcitabine predictor. We show the distribution of their weights annotated with the ten largest weights (resistance) and ten smallest weights (sensitivity). (**D**) Interpretation of Paclitaxel predictor. (*D*.*1*.) We display the positions of genes associated to two significantly enriched gene sets, both to resistance: “Basaki YBX1 target UP” and “constitutive signaling by PI3K-aberrant signaling”. (*D*.*2*.) Interaction terms for Paclitaxel, that make 12% of the predictor, alongside the top-10 resistant and top-10 sensitive interactions.

Cytotoxic drugs such as Gemcitabine or Paclitaxel have complex modes of actions involving different pathways, crosstalk between which remains challenging to understand. Since the predictions of these two drugs showed a significant association in both PDXs and patients, we set out to interpret the mechanisms of sensitivity or resistance inferred by our predictor. In Gemcitabine (**Supp. Table 3**), we observe that over-expression of the CDC42 pathway is a significant marker of resistance (FDR = 0.012, **Figure 5**C.1.) together with pathways linked to microtubule formation and cell migration (**Supp Figure 13**), both known to be promoted by CDC42^45^. Together, these enriched pathways highlight CDC42 over-expression as a potential mechanism of Gemcitabine resistance, which suggests the use of CDC42 inhibitors^46,47^ for Gemcitabine-resistant tumors. Another interesting finding is the significant enrichment of TNF*α* signaling in the sensitive portion (FDR=0.046) (**Figure 5**C.1.). A clinical trial has shown that co-administration of TNF with gemcitabine improves patient survival and further inhibits tumor growth^48^, lending additional credence to this finding. Last, we observe a concentration of sensitive interactions involving BLK, a pro-apoptotic Src-proto oncogene involved in B-cell signaling and differentiation (**Figure 5**C.2.). Since hematopoietic cell lines respond better to Gemcitabine, these interactions can either act as a tissue-type marker or could potentially represent a sensitive pathway. Finally, we looked for enriched pathways in Paclitaxel predictor (**Figure 5**D.1., **Supp. Table 4**) and observed three potential mechanisms of resistance. We first observe that the resistant linear coefficients are significantly enriched in genes linked to silencing of YBX1^49^ (FDR=0.106), a gene associated with proliferation in certain tumor types^50^. In ovarian cancer, YBX1 has been shown to regulate ABCB1 expression levels, a gene related to Paclitaxel resistance^52–56^. Our pan-cancer analysis therefore further supports the role of drug transporters in Paclitaxel resistance. Second, we observe a significant resistant enrichment for PI3K activation (FDR=0.18), which is corroborated by the observed activation of PI3K/AKT/mTOR signaling pathway in Paclitaxel-resistant cancer cells^56,57^. Moreover, a recent investigation suggests that PI3K catalytic subunits can regulate ABCB1 expression^58^. Finally, when it comes to the non-linear part, we observe a concentration of Fibroblast growth factors interactions in the resistant parts of non-linearities, in particular FGF3, FGF20 and FGF8 and FGF4 (**Figure 5**D.2., **Supp. Table 4)**. This behavior, although suggested by previous studies^59,60^, is all the more interesting as cell lines do not contain any micro-environment that would elicit such resistance.

## Discussion

We introduced an approach to integrate pre-clinical and clinical data in a fully unsupervised way. Our approach geometrically aligns sample-to-sample similarity matrices and extracts directions of important variations for both datasets, without requiring any sample-level pairing. By performing a geometrical alignment instead of a direct distribution comparison, our approach limits the effect of a potential sample selection bias. This geometrical alignment is implicitly performed in a space induced by our similarity function, which enables the integration of various non-linearities, corresponding to hypothesis made on the system. Although we restricted ourselves to a Gaussian similarity, designing similarity functions that incorporate a wide range of prior knowledge is a potentially promising avenue. Learning the similarity matrix, e.g. using multiple kernel learning^61^ or deep learning methods such as variational auto-encoders^62^, may also help increase performance.

TRANSACT compares directions computed using Kernel PCA, but our approach can be extended to other basis expansion methods by modulating the way the coefficients *α*^*s*^ and *α*^*t*^ are computed. More generally, our method is versatile, generalizable, and can be applied beyond the scope of our study, e.g. to integrate single cell sequencing data.

We showed that the consensus features can be used to build translatable predictors of drug response. Although we do not require a strong covariate shift assumption as in a previous study^63^, we do assume the functions modeling the response from these consensus features follow the same monotonicity in pre-clinical models and human tumors. This assumption, albeit reasonable, may be questioned.

In this study, we limited ourselves to gene expression. Making use of other genomics levels – e.g. mutations, copy number, methylation, chromatin accessibility – may help refine the prediction by providing additional signal. The integration of our approach with multi-omics integration strategies^64,65^ may offer a solution to the translation of multi-omics signatures.

Finally, we were able to predict response in patients that received a particular treatment, either as monotherapy or in combination with other therapies, even though the cell line predictors were trained on monotherapies only. We envision that large-scale, cell line combination screens that take potential synergistic effects between drugs into account will provide training data for the next generation of cell line predictors to be transferred to human tumors in order to further enhance prediction of patient response. Moreover, in this study, we focus on cytotoxic and targeted therapies, still widely used in the clinic. The recent advent of immuno-therapies calls for methods with the ability to predict the clinical response from model systems. This requires model systems capable of mimicking the action of the immune system and screening technologies able to measure the response for large panels. We believe that our approach can be extended to such problems once data is made available.

## Methods

### Public data download and pre-processing

#### GDSC dataset, download and processing

We made use of the GDSC1000 cell line panel^14^, which contains complete molecular profiles for 1,049 cell lines (**Supp Figure 2**). Gene expression is provided in the form of both read counts and FPKM. For both settings, we corrected the dataset for library-size using TMM^66^ and log-transformed the corrected read counts^67,68^. Finally, we performed a gene-level mean-centering and standardization.

#### Novartis PDXE dataset, download and processing

We made use of NIBR PDXE dataset for patient-derived xenografts^15^, which contains the gene expression profiles of 399 PDXs (**Supp Figure 3**). Gene expression is provided in the form of FPKM. We corrected for library-size using TMM^66^ and log-transformed the corrected read counts^67,68^. Finally, we performed a gene-level mean-centering and standardization.

#### TCGA dataset, download and processing

We made use of the TCGA dataset for analyzing human biopsies^69^, which comprises 10,347 human tumors (**Supp Figure 4**). Gene expression is provided in the forms of both read counts and FPKM and we used the same pre-processing pipeline as for GDSC. Response data have been obtained from *Ding et al*^70^. Following *Ding et al*, for each drug, we consider the response to patients who were administered a particular drug either as monotherapy or in combination with other drugs.

### Hartwig Medical Foundation dataset (HMF) download and processing

We validated our approach on a cohort of 1,049 patients provided by the Hartwig Medical Foundation – referred to as *HMF* (**Supp Figure 5**A). Gene expression was measured for each metastatic sample prior to indicated drug regimen. We used MultiQC for quality control^71^, salmon v1.0.0 for alignment to reference transcriptome^72^, and finally edgeR for gene-level quantification^73^. Comparison with results obtained using STAR^74^ and featureCounts^75^ shows high degree of concordance (**Supp Figure 5**D) and we used this comparison to refine our filtering. Read counts were then processed using the same pipeline as in GDSC and TCGA.

Drug response was measured in 802 unique metastatic samples using the RECIST criteria. Response was measured differently for each patient (**Supp Figure 5**B) with most patients having one single measure of response around 10 to 15 weeks after treatment started (**Supp Figure 5**C). Since we are interested in the response of the drug given the molecular characterization measured, we considered for each patient the first response after treatment. Since most drugs are administered in combination, we considered, for each drug, all the patients that received it, whether in combination with other drugs or as monotherapy. For instance, in the case of Gemcitabine, we predicted drug response for all patients that received Gemcitabine as part of their treatment strategies.

### Measure of drug response

In our different analysis, we rely on drug response measurements, either to train a predictor (GDSC), or to validate it (PDXE, TCGA and HMF). For cell lines (GDSC), drug response is measured as *Area Under the drug response Curve*, referred to as AUC. For PDX, drug response is measured as *Best Average Response*, which corresponds to the relative variation of tumor volume induced by a certain treatment. For both AUC and Best Average Response, large values are associated with resistance. For TCGA and HMF, clinical responses have been measured using the RECIST criteria^76^. Based on various metrics, patients get assigned to one of the following four groups: *Complete Response* (CR), *Partial Response* (PR), *Stable Disease* (SD) and *Progressive Disease* (PD). Following the division used in previous works^21,23^, we divide TCGA patients in two categories: *Responders* (CR and PR) and *Non Responders* (SD and PD). For HMF, we discriminate between each possible couple: PR vs PD, PR vs SD and SD vs PD. Since only a couple of patients showed a complete response, we did not consider these patients.

### Mathematical settings

We denote by *p* the number of genes. We consider one source dataset 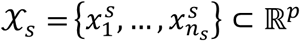 and one target dataset 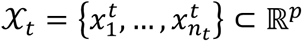 with corresponding source and target data matrices 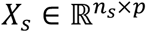 and 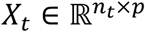.

We consider a similarity function *K* -- also called *kernel function* -- that assigns to two samples a scalar value that is large for similar samples and small for dissimilar samples. In this work, we assume the kernel to be positive semi-definite (p.s.d.), which implies^77^ that there exists a Hilbert space *ℋ* and a mapping *φ*: ℝ^+^ ↦ *ℋ* such that

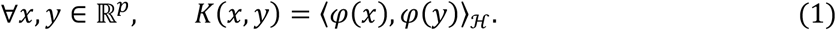

In particular, we use two kernels:

▪ Linear kernel: *K*^*linear*^(*x, y*) = *x*^*T*^*y*.
▪ Radial Basis Function, also referred to as “Gaussian”: 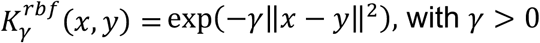.

We denote by *K*_*s*_ the matrix of similarity between source samples, *K*_*t*_ between target samples and *K*_*st*_ the matrix of similarity between source and target, formally:

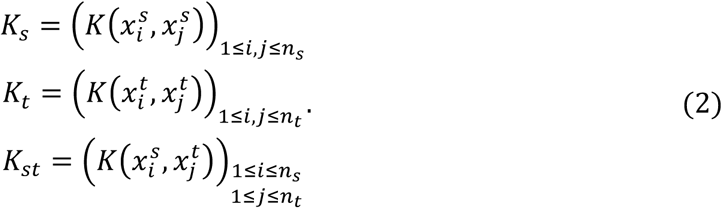

### Implicit mean centering

In a linear-setting, it is standard to perform a gene-level centering prior to any statistical analysis. In the case of non-linear analysis, we perform a feature-level centering in the embedding space *ℋ* implicitly using Equation (1). We perform this implicit centering independently in source and target datasets. As shown in (Subsection Supp. 2), the corresponding kernel matrices are:

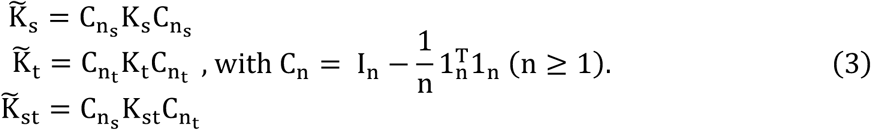

### Kernel PCA by eigen-decomposition of centered kernel matrix for capturing directions of principal variance

Using the embedding introduced in Equation (1), the similarity matrices from Equation (3) can be seen as sample-covariance matrices and therefore decomposed to compute principal components inside the embedded space *ℋ*, a procedure known as Kernel PCA^36^. We perform Kernel PCA on source and target data independently to compute *d*_*s*_ and *d*_*t*_ principal components respectively. Kernel PCA on the source dataset consists of an eigen-decomposition of the matrix 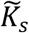, yielding 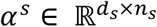, while Kernel PCA on the target dataset decomposes 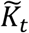, yielding 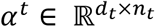 (Def Supp 3.1).

### Comparing and aligning pre-clinical and tumor non-linear principal components

Similarly to the “cosine similarity matrix” in other related works^24,28^, we define the *non-linear cosine similarity matrix* ***M***^*K*^ as the matrix that geometrically compares the source NLPCs to the target NLPCs (Def Supp 5.1). This matrix can be computed as follow (Prop Supp 5.2):

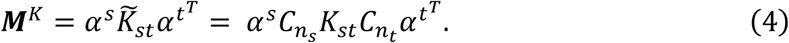

In a first step of our domain adaptation approach, we use the matrix ***M***^*K*^ to align NLPCs, yielding *non-linear principal vectors s*_1_,. ., *s*_*d*_ for the source and *t*_1_,. ., *t*_*d*_ for the target domains, with *d* = min(*d*_*s*_, *d*_*t*_) (Def Supp. 4.1). These principal vectors are pairs of vectors: one linear combination of source NLPCs and one linear combination of target NLPCs, ordered by decreasing similarity with the first pair being the most similar. The computation of these PVs relies on the Singular Value Decomposition^78^of 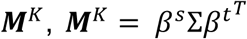 that helps us define the source and target *sample importance loadings ρ*^*s*^ and *ρ*^*t*^ as follows (Def. Supp. 5.4)

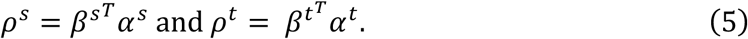

We also define the principal angles *θ*_1_,. ., *θ* _*d*_ as follows (Def. Supp. 5.6)

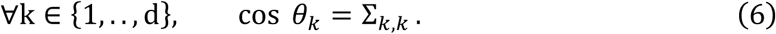

Owing to the duality elicited in Equation (1), these principal vectors can be seen both as functions that map samples in cell-view to a one-dimensional vector, or a vector onto which we project the sample embedding (Proposition Supp. 5.8).

### Interpolation between kernel principal vectors for balancing effect of source and target

Each pair of principal vectors contains two vectors that are geometrically similar. Projection on them will create two highly correlated covariates that would not be optimal for subsequent statistical treatment. In order to compute one vector out of each pair, we interpolate between the source and the target PV within each pair (Def Supp 6.2). For the *k*^*th*^ PV, the interpolation is modulated by a coefficient *τ*_*k*_ that ranges between 0, when the interpolation returns the source PV, and 1, when the interpolation returns the target PV. This interpolation between vectors within each PV pair relies on two functions Γ(*τ*) = [Γ_1_(*τ*_1_),. ., Γ _*d*_(*τ* _*d*_)]^*T*^ and ξ(*τ*) = [ξ_1_(*τ*_1_),. ., ξ_*d*_(*τ*_=_)]^*T*^ defined as (Definition Supp. 6.1):

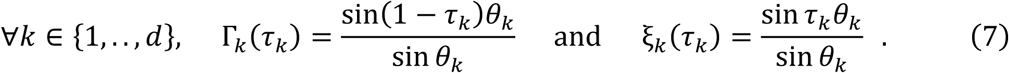

For a set of *d* interpolation coefficients [*τ*_1_,. ., *τ* _*d*_], we compute the projection of source and target datasets 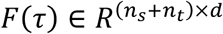 as follows (Theorem Supp. 6.6)

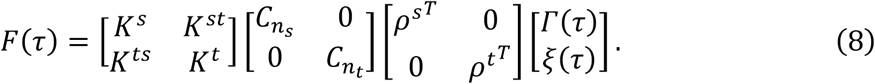

Such an interpolation between PVs balances the effect of source and target datasets. We prove that, in the case of a linear kernel, our interpolation scheme is equivalent to the one from previous approaches^26,79^ (Supp Subsection 8).

Within each pair of PVs, we select one intermediate representation where the source and target projections match the most. For the *k*^*th*^ PV-pair, we compare the source and target projected data using a Kolmogorov-Smirnov statistic and select the interpolation coefficient 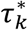 where the statistic is minimal. We obtain a set of optimal interpolation coefficients *τ*^*^ ∈ [0,1]^*d*^ when, for each PV, source and target influence are balanced. We call the corresponding vector *consensus features*. These consensus features show the minimal difference between source and target domain, a theoretical necessary condition for domain adaptation^80^.

### Prediction using Elastic Net

In order to predict drug response, we use Elastic Net regression^81^. Elastic Net is a linear model that imposes two penalties on the coefficients to predict: an *ℓ*_)_penalty that leads to a sparse model and an *ℓ*_2_ penalty that jointly shrinks correlated features. We chose Elastic Net first because it has repeatedly been shown in the drug response prediction literature to give equivalent, if not better, predictive performance compared to complex models^16,18,38^. Using a linear classifier limits the complexity and therefore makes the transfer more robust in practice.

### Taylor expansion of the similarity function for interpretability of the model

In the case of RBF, we perform a PCA in an infinite-dimensional feature space. Although this space cannot be analytically computed, it can be approximated using a Taylor expansion^82^ (Subsection Supp. 7). For the *q*th consensus feature, we differentiate three kinds of contributions (Def Supp. 7.4):

- *Offset 𝒪*_*q*_: a Gaussian term that models the squared depth. For each sample *x* ∈ *R*^*p*^, it corresponds to exp (−*γ* ‖*x*‖^2^).
- *Linear contributions* 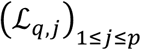: a linear term, resembling the expression of one gene. For each sample *x* ∈ *R*^*p*^ and gene *j* ∈ {1,. ., *p*}, it corresponds to *x*_*j*_exp (−*γ* ‖*x*‖^2^).
- *Interaction terms* 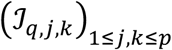: an interaction term that expresses the product of two genes. For each sample *x* ∈ *R*^*p*^ and gene *j, k* ∈ {1,. ., *p*}, it corresponds to *x*_*j*_*x*_*k*_exp (−*γ* ‖*x*‖^2^).

These contributions can be computed from sample importance loadings of consensus features (Prop. Supp. 7.7). We consider the contributions’ sum-of-squares as a geometrical proportion since these sum up to one (Def. Supp. 7.8).

In order to look for enrichment in a particular consensus feature, we look for enrichment of particular gene sets^83^. Specifically, for the linear contribution, we compute the loading of all linear terms (Equation Supp. 48) corresponding each to one gene and we performed a Pre-Ranked gene set enrichment analysis with FDR correction at 20% and 1000 permutations. Since these loadings correspond to a Euclidean geometric proportion, we used a squared statistic to compare them.

### Comparison to competing approaches

We compare TRANSACT to four different approaches. The two first approaches consist in applying a regression model trained on a source dataset to a target dataset without any correction; we use one linear Elastic Net model (referred to as *Elastic Net*) and a non-linear neural network model (referred to as *Deep Learning*). In both cases, we perform a grid-search 5-fold cross-validation on cell lines to select the model with the best performance: on *ElasticNet* we vary the *ℓ*_1_ ratio and the total regularization ; on *Deep Learning*, following the protocol from *Sakellaropoulos et al*^21^, we use an hyperbolic-tangent activation function while varying the global network structure, the *ℓ*_2_ penalty and the input and output dropout levels (**Supp Table** 5).

The other two approaches first correct the signal and then train a regression model. The third approach (referred to as *ComBat + Deep Learning*) reproduces the approach from *Sakellaropoulos et al*^21^ by first performing a ComBat technical batch effect correction between source and target, and then applying a neural network on the corrected signal, similar to *Deep Learning* (**Supp Table** 6). The last competing approach, referred to as *PRECISE*, consists in using a linear similarity function followed by an Elastic Net model, which is equivalent to PRECISE (**Supp Material**). For the two deep learning approaches, we first performed cross-validation on the source dataset (with or without correction) to select the hyper-parameters and the network structures with the largest predictive performance. We then re-initialize the network and train it on the complete GDSC dataset.

### Availability of data and materials

TRANSACT is available as a Python 3.6 module (https://github.com/NKI-CCB/TRANSACT). All our experiments are reproducible and use state-of-the-art libraries^84–89^ (https://github.com/NKI-CCB/TRANSACT_manuscript). The dataset(s) supporting the conclusions of this article are available in the “transact_manuscript” repository of the aforementioned GitHub page, except for the HMF data, which can only be obtained upon request to the Hartwig Medical Foundation.

## Supporting information

Algorithm Derivation

## Acknowledgment

This publication and the underlying study have been made possible partly on the basis of the data that Hartwig Medical Foundation and the Center of Personalised Cancer Treatment (CPCT) have made available to the study.

We thank Mirrelijn van Nee (VUmc), Sander Canisius (NKI), Osman Kayhan (TU Delft), Tycho Bismeijer (NKI), Stavros Makrodimitris (TU Delft), Tesa Severson (NKI), Joe Siefert (NKI), Joris van de Haar (NKI), Ahmed Mahfouz (LUMC), Tamim Abdelaal (TU Delft), David Tax (TU Delft), Bram Thijssen (NKI) and Ziqi Wang (TU Delft) for useful discussions. We thank Mohammed Charrout (TU Delft), Marie Corradi (NKI), Lisa Koob (NKI) and Wouter Kouw (TU Eindhoven) for careful reading of the manuscript.

This work was supported by ZonMw TOP grant COMPUTE CANCER [40-00812-98-16012].

## Authors contribution

S.M., L.F.A.W., M.J.T.R., M.L. and M.A.vd.W. designed the study. S.M. performed the experiments. L.F.A.W., M.J.T.R. and M.L. supervised the experiments. S.M., L.F.A.W., M.J.T.R., M.L. and M.A.vd.W. analyzed the results. S.M. and M.L. developed the mathematical framework. S.M. developed the software package. S.M. and D.J.V. aligned the HMF data. K.M. and A.G. interpreted the GSEA results. S.M., L.F.A.W., M.J.T.R. and M.L. wrote the manuscript. All authors read and approved the manuscript.

---

**Supplementary Figure 1.**
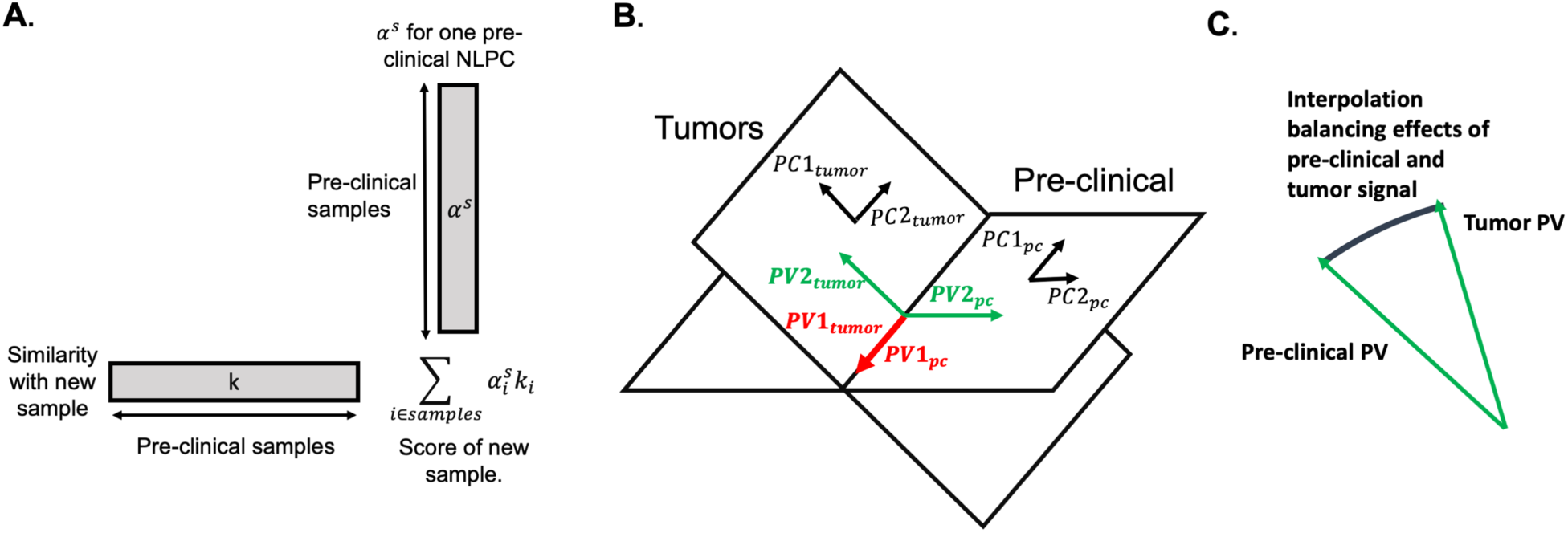
<u>Visual explanation of geometric alignment</u>. **A**: Difference between importance scores (*α*^*s*^, *α*^*t*^) and projected scores. Since the space induced by the similarity function *K* is intractable, we use a dual representation of the NLPC in terms of samples: the importance scores. To project samples on NLPCs, one needs to compute the similarity between this sample and all of the samples used to gauge the NLPC. The projected score is obtained by taking the vector-product between this similarity vector and the importance scores. The same rational holds principal vectors that are represented by *γ* ^*s*^and *γ* ^*t*^. **B**: Visual example of principal vectors (PV). We here consider 3 genes (features) and 2 NLPCs. The pre-clinical (source) and tumor (target) NLPCs intersect in one direction, which form the pair of closest vectors: the first PV forms the pair of the two red vectors – although these are identical. The second pair of PVs is defined orthogonally to the red pair. This defines the green vectors (with a swap in direction for visual purposes). These pairs reconstruct the original NLPC spaces and are ordered by similarity. **C**: Interpolation between PVs. For one pair of PVs – e.g. the green one in B – source and target vectors are different. In order to generate one robust vector out of these two and avoid redundancy, we draw an arc between these two vectors. We then project source and target datasets onto these interpolated vectors and select one intermediate representation where source and target projected signals are maximally matched. This optimal intermediate vector is called the consensus feature.

**Supplementary Figure 2.**
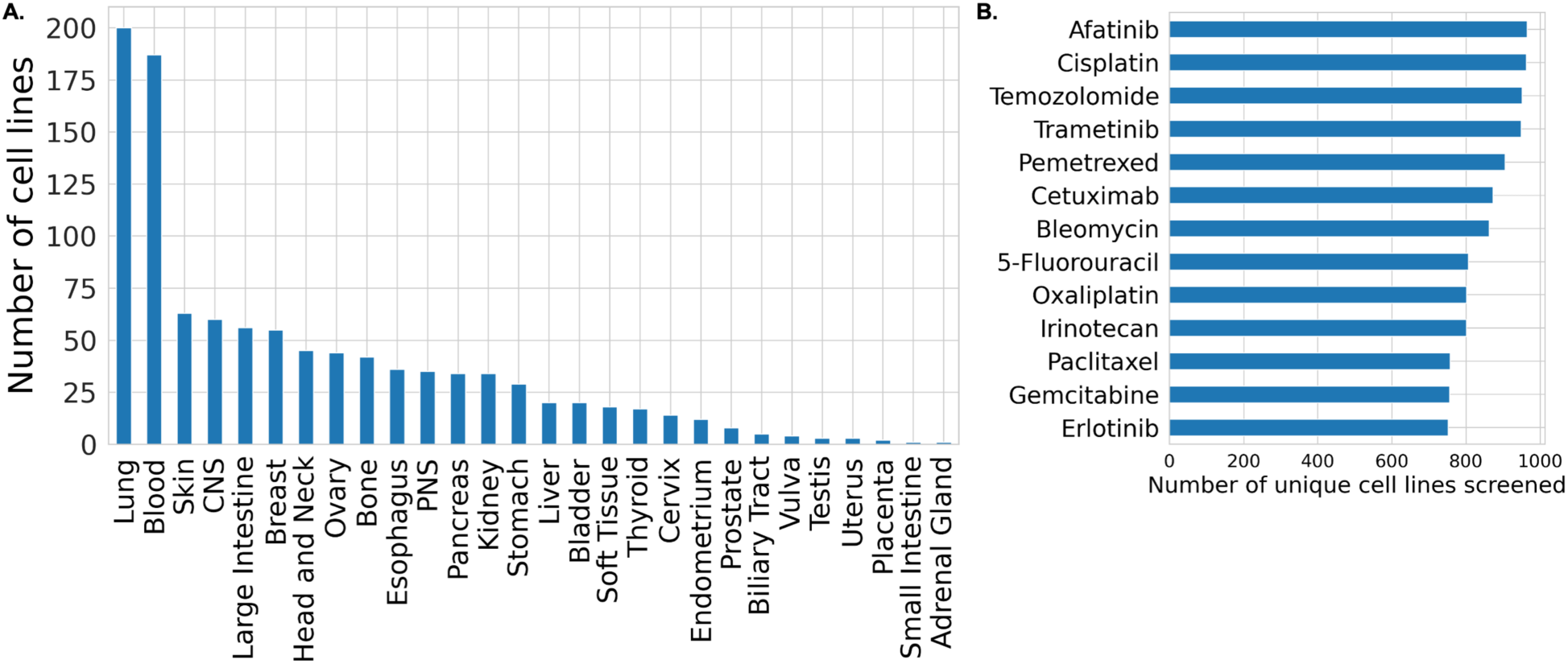
<u>Composition of the GDSC dataset (cell lines).</u> We make use of the GDSC1000 cell line panel^14^. **A**: Number of cell lines per tissue type. **B**: Number of cell lines screened for each drug that we used in our experiments.

**Supplementary Figure 3.**
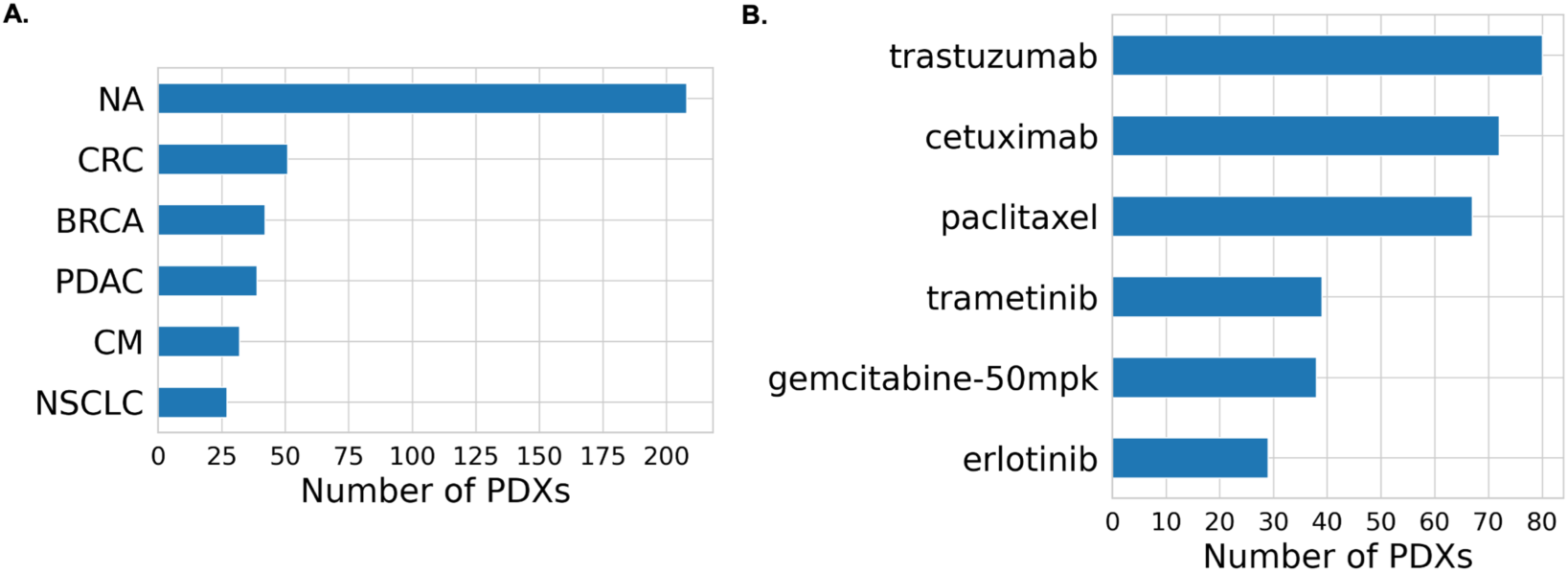
<u>Composition of the NIBR PDXE dataset (patient derived xenografts).</u> We make use of the NIBR PDXE patient derived xenograft panel^15^. **A**: Number of PDXs per tissue type. **B**: Number of unique PDXs screened for each drug that we used in our experiments.

**Supplementary Figure 4.**
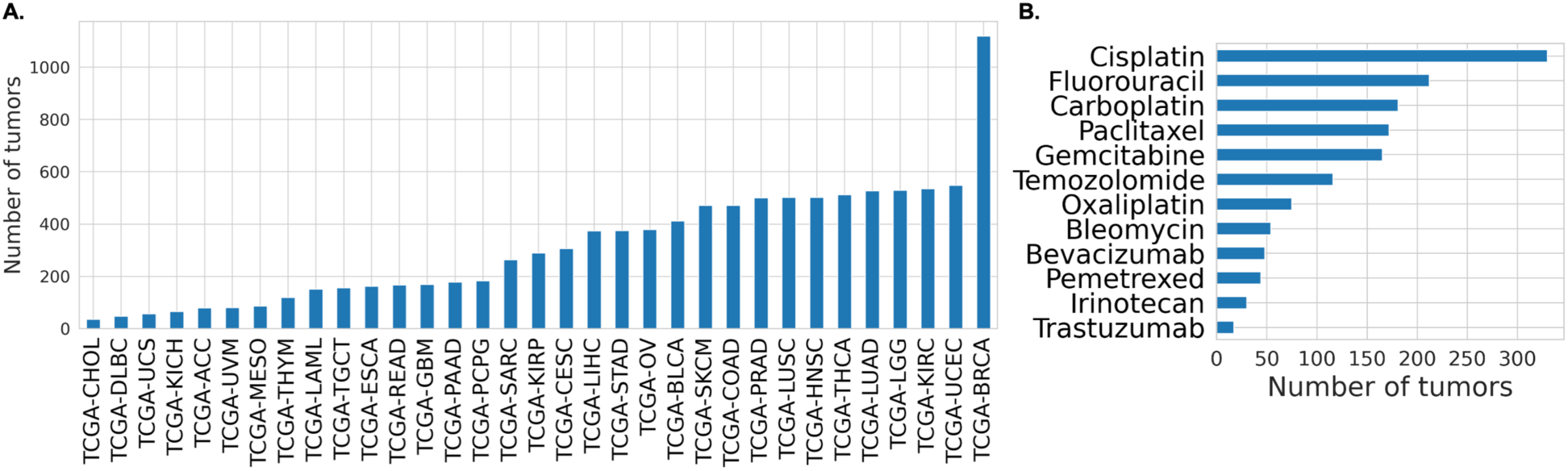
<u>Structure of the TCGA dataset (primary tumors)</u>. We make use of the TCGA dataset for primary tumors. **A**: Number of samples per cancer type. **B**: For each drug, number of samples with known response.

**Supplementary Figure 5.**
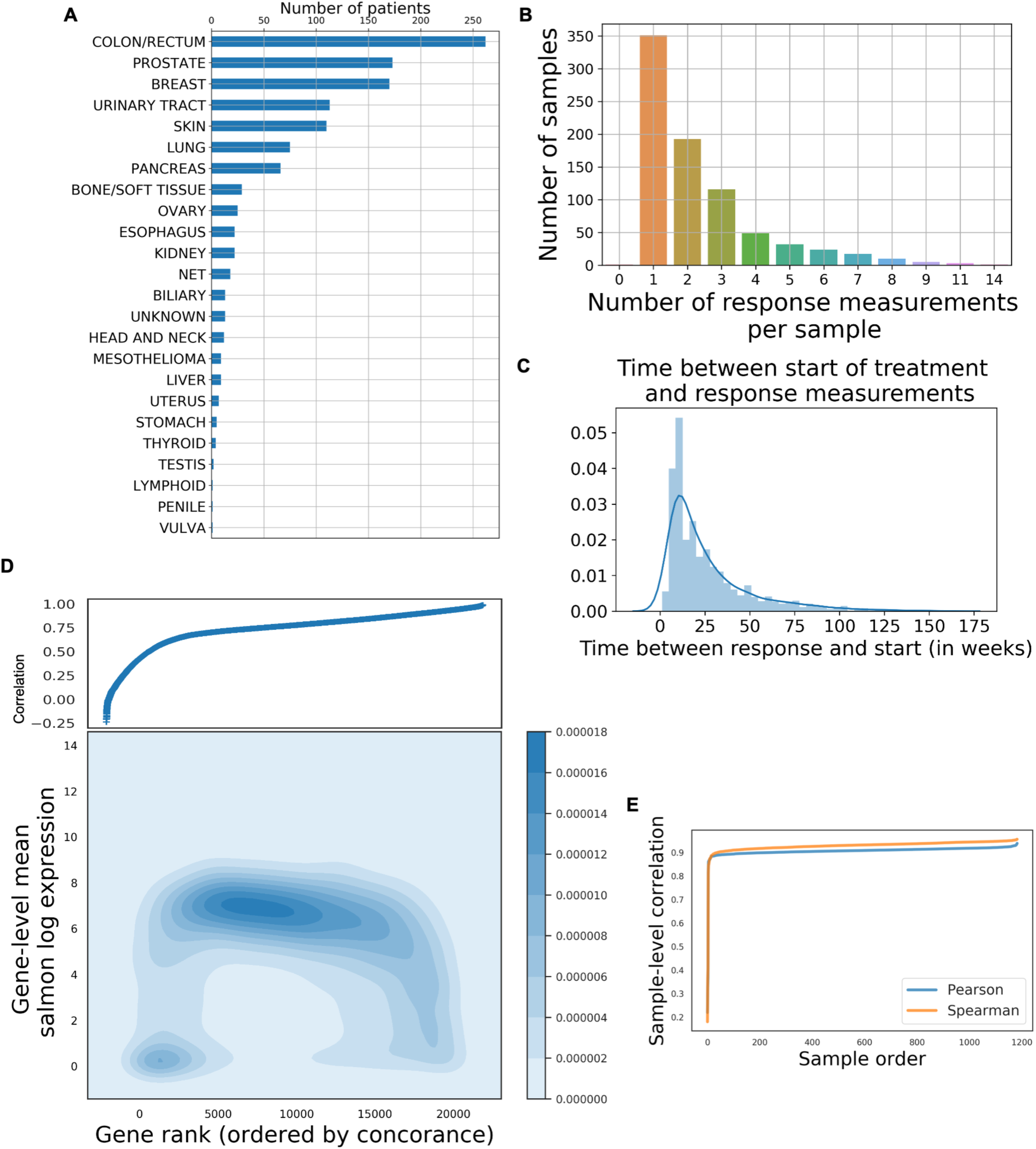
<u>Structure of the HMF dataset (metastatic lesions).</u> We make use of the Hartwig Medical Foundation (HMF) dataset for metastatic lesions. (**A**) Number of samples per cancer type (primary tumor location). (**B**) For each patient, number of response measurements made. For further analysis, we considered the first response measure – i.e. first measure after treatment start. (**C**) Histogram of number of weeks between treatment start and response measurement. (**D**) For each protein coding gene, we measure the Spearman correlation between read counts obtained using Salmon and STAR alignment tools using all samples in the HMF dataset. We then ranked genes based on the obtained Spearman correlation (x-axis) and plotted it against the mean-expression of these genes obtained using Salmon (y-axis). Since lowly concordant genes tend to have low expression, we put a threshold at *corr* = 0.5 and discarded genes below this threshold. (**E**) After the previous selection, we computed the sample-level Pearson and Spearman correlations between read counts obtained with STAR and Salmon. All samples but five show a correlation above 0.8 – these were discarded. We finally further restricted to genes from the mini-cancer genome.

**Supplementary Figure 6.**
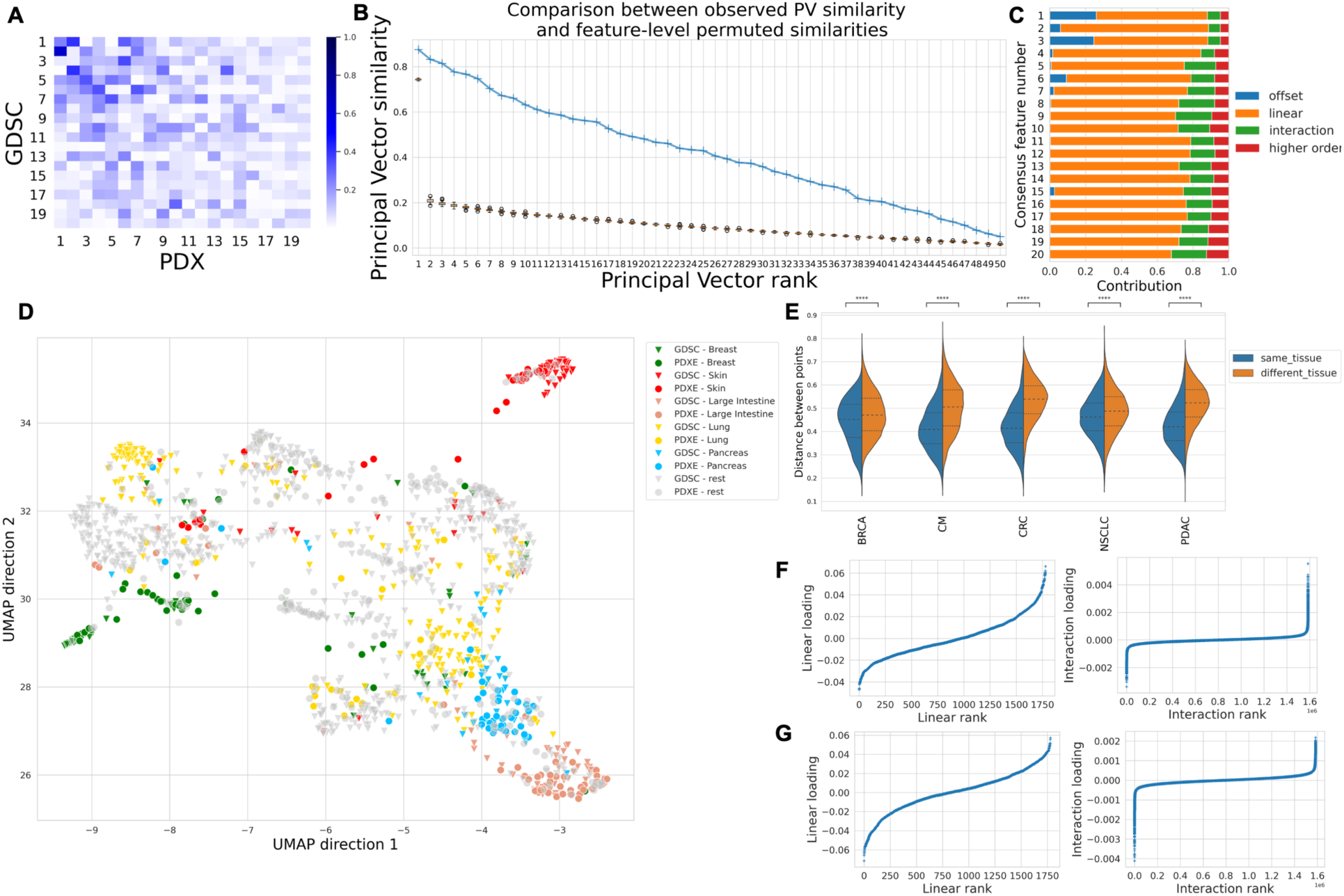
<u>Analysis of consensus features between cell lines (GDSC) and PDXs with</u> *<u>γ</u>* <u>= 0.0005.</u> We use a Gaussian similarity matrix with hyper-parameter *γ* = 0.0005 and run TRANSACT. (**A**) Cosine similarity between the 20 top source and target NLPCs. (**B**) Similarity between principal vectors (blue line) alongside the similarity obtained after gene-level permutation on GDSC (boxplots). (**C**) For each consensus feature, proportion of offset, linear and interaction term. (**D**) UMAP of data projected on the consensus features, colored by tissue of origin. (**E**) For each tissue type in PDXs, we compare the distances between corresponding PDXs with cell lines from the same tissue of origin (blue), or from another tissue (orange). (**F**) For the first consensus feature, sorted contribution of each linear features (i.e. gene, left) and interaction terms (right). (**G**) For the second consensus feature, sorted contribution of each linear features (i.e. gene, left) and interaction terms (right).

**Supplementary Figure 7.**
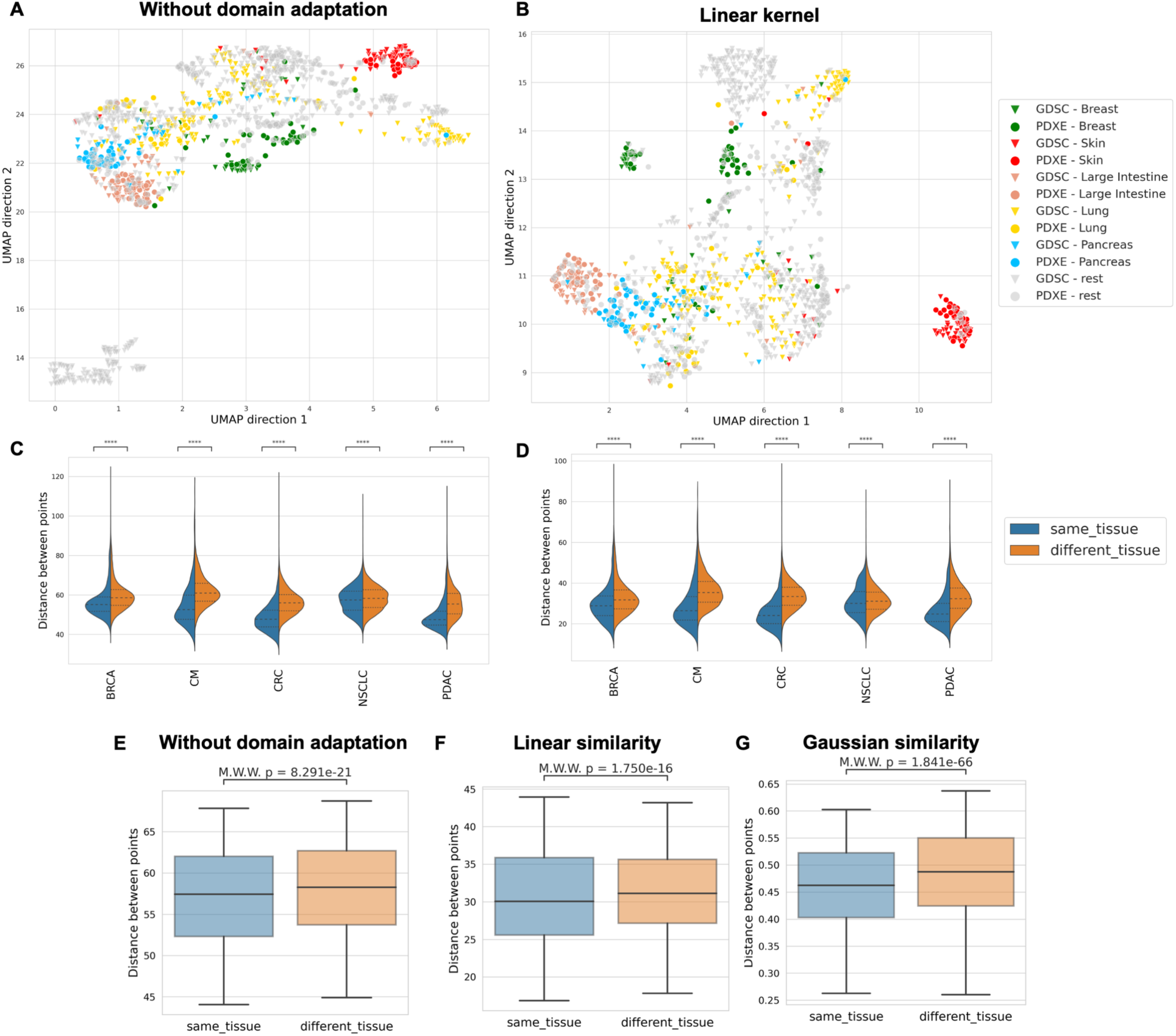
<u>Tissue clustering without domain adaptation and with PRECISE alignment between GDSC and PDXE</u>. (**A**) UMAP plot of cell lines and PDXs colored by tissue type without any domain-adaptation. Data was normalized prior to performing UMAP: cell lines and PDXs were independently mean-centered and scaled to unit variance. (**B**) UMAP plot of cell lines and PDXs colored by tissue type after projection on consensus features obtained with linear PRECISE. (**C**) Comparison of distances between PDXs and cell lines from the same tissue type (blue) or from a different tissue type (orange) without domain adaptation. (**D**) Comparison of distances when using linear PRECISE. We zoom in on lung (NSCLC) without domain adaptation (**E**), with linear PRECISE (**F**) or with TRANSACT (**G**) using same setting as in **Supp Figure 6**.

**Supplementary Figure 8.**
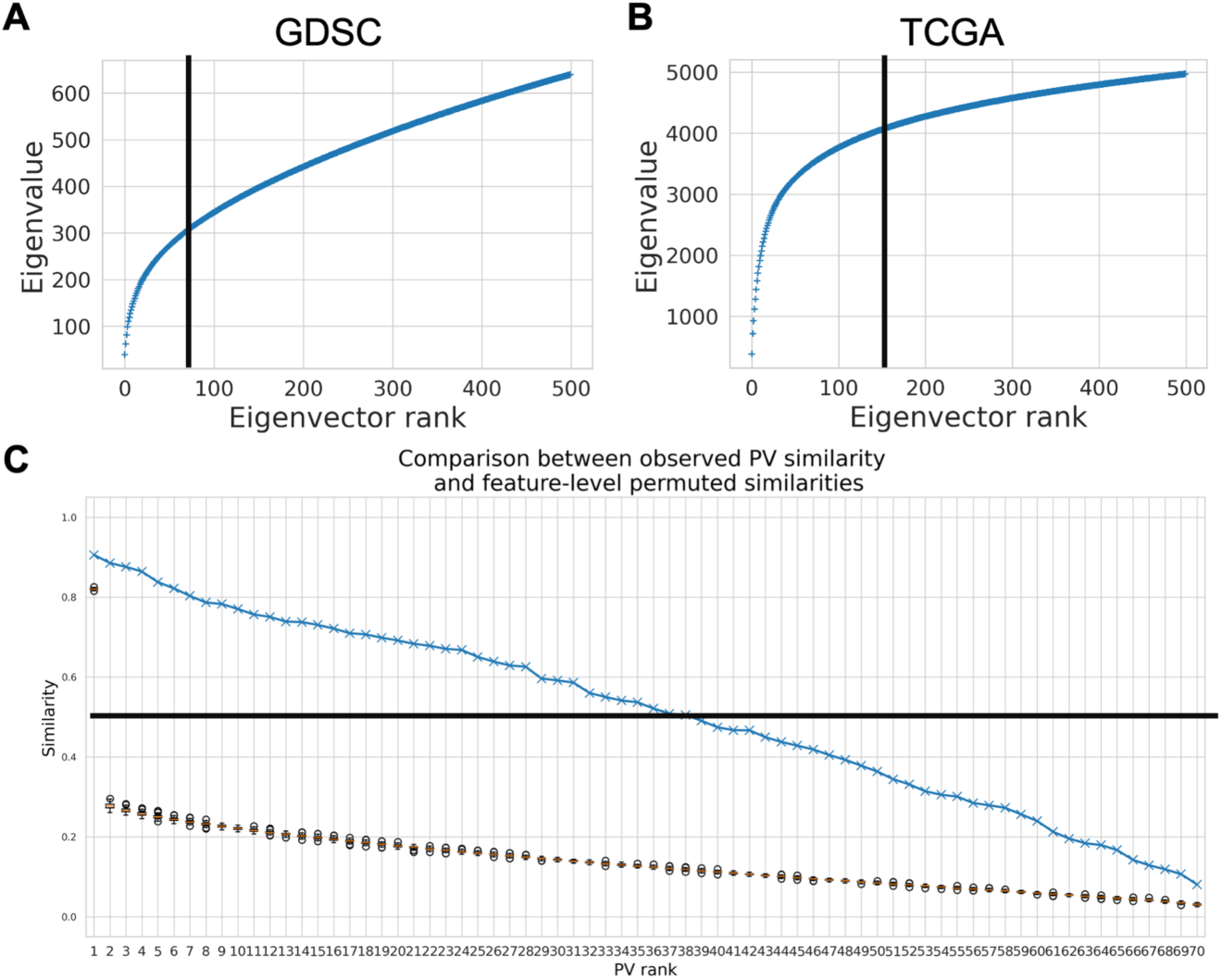
<u>Choice of the number of NLPCs and consensus features between GDSC and TCGA.</u> (**A**) Cumulative sum of eigenvalues of 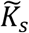 (GDSC) with *γ* ^*^ = 5 × 10^−4^. The cumulative sum increases steeply, reaches an inflexion points and then follows an almost-linear behavior. We select all the NLPCs before this almost-linear zone, corresponding to 75 NLPCs. (**B**) Cumulative sum of eigenvalues of 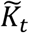 (TCGA) with *γ* ^*^ = 5 × 10^−4^. Following similar reasoning as in (**A**), we restrict the study to the first 150 NLPCs. (**C**) Similarity between PVs when 75 NLPCs are considered for GDSC and 150 for TCGA. We observe that the 33 first PVs have a similarity above 0.5 (our cut-off) and round the selection to 30 PVs.

**Supplementary Figure 9.**
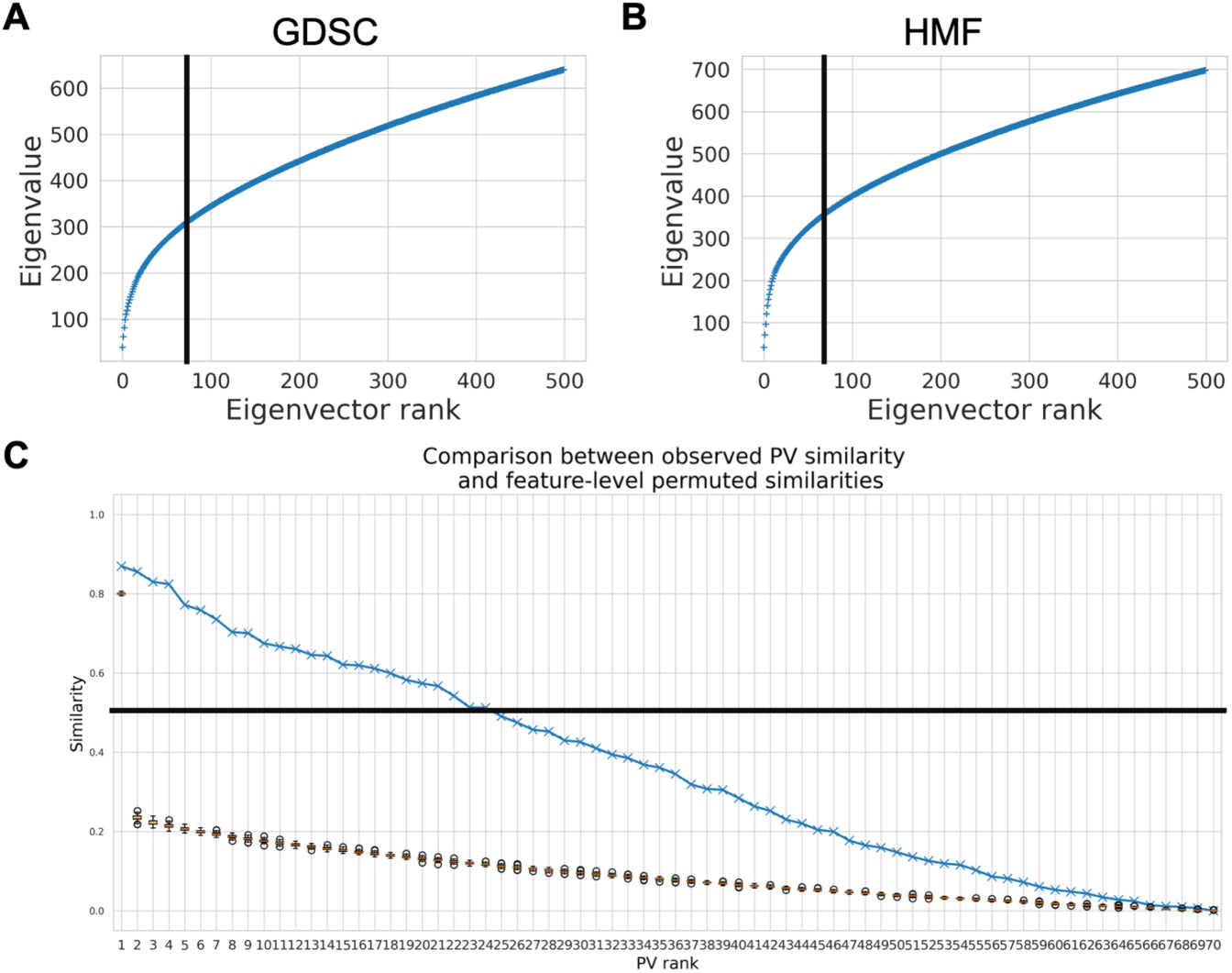
<u>Choice of the number of NLPCs and consensus features between GDSC and HMF.</u> (**A**) Cumulative sum of eigenvalues of 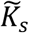 (GDSC) with *γ* ^*^ = 5 × 10^−4^. The cumulative sum increases steeply, reaches an inflexion points and then follows an almost-linear behavior. We select all the NLPCs before this almost-linear zone, corresponding to 75 NLPCs. (**B**) Cumulative sum of eigenvalues of 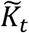 (HMF) with *γ* ^*^ = 5 × 10^−4^. Following similar reasoning as in (**A**), we restrict the study to the first 75 NLPCs. (**C**) Similarity between PVs when 75 NLPCs are considered for both GDSC and HMF. We observe that the 21 first PVs have a similarity above 0.5 (our cut-off) and round the selection to 20 PVs.

**Supplementary Figure 10.**
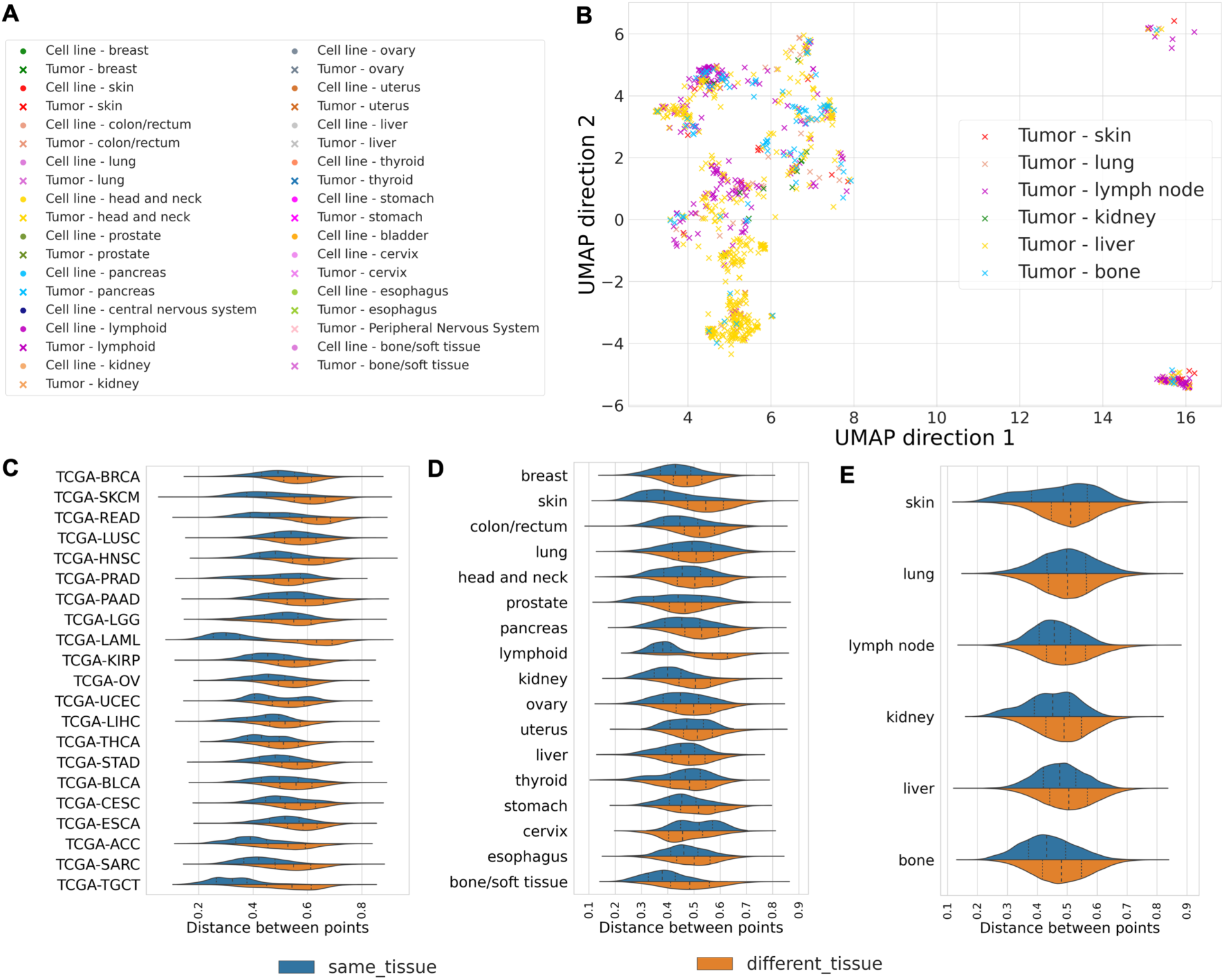
<u>Pan-cancer consensus features between cell lines and tumors conserve tissue type information (Supplement of Figure 3)</u> **A**: Legend of UMAP plots for **Figure 3**C-D. **B**: UMAP plot of HMF metastatic lesions (same as Figure 3D) colored by metastatic site. **C**: In TCGA, for each tumor type, distance between tumors and cell lines from similar (blue) and non-similar (orange) tissue. **D**: In HMF, for each primary tumor type, distance between metastatic sample and cell line from similar and non-similar tissue of origin. **E**: In HMF, for each metastatic site, distance between metastatic sample and cell line from tissue of origin similar (blue) or dissimilar from the metastatic site.

**Supplementary Figure 11.**
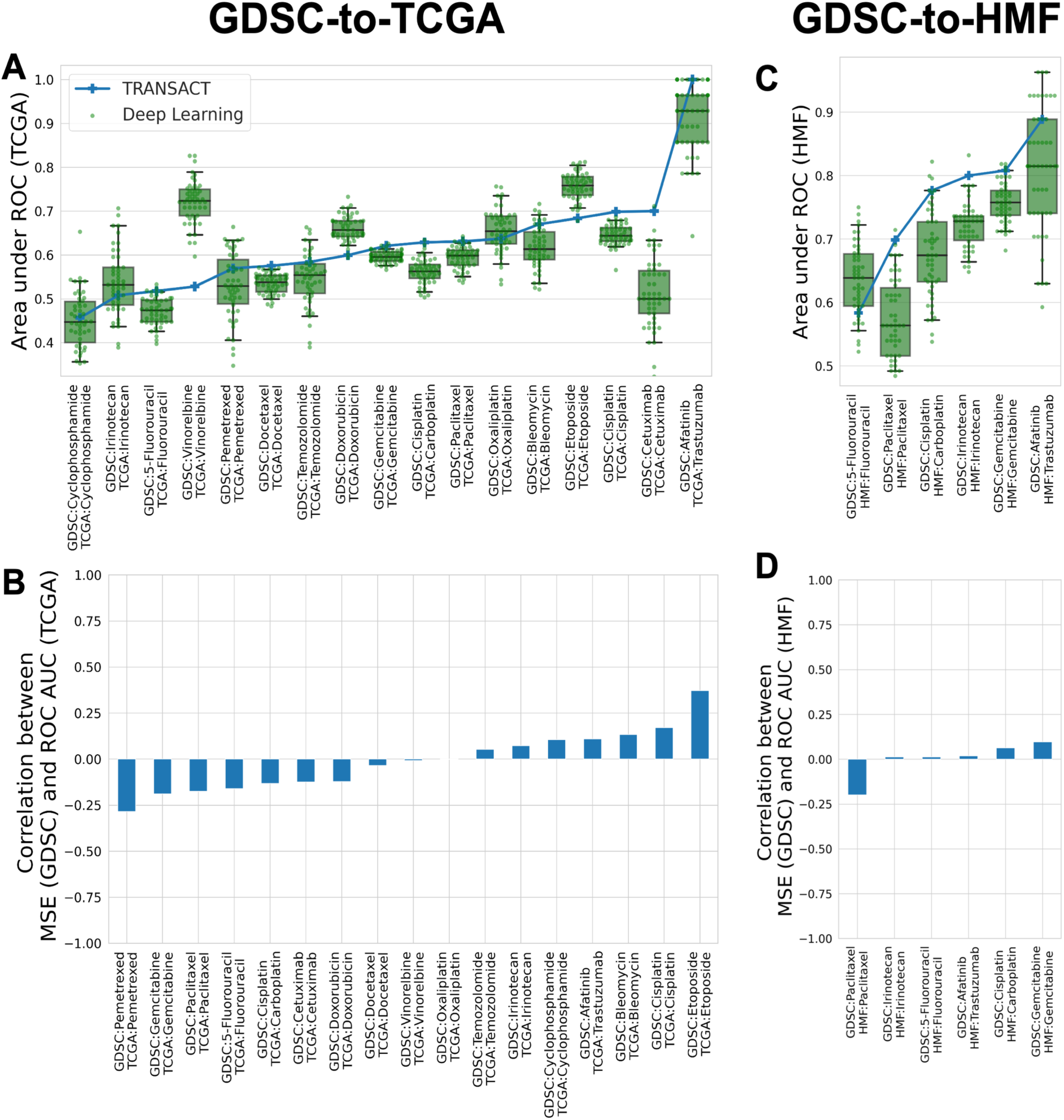
<u>Impact of initialization on results for the</u> *<u>Deep Learning</u>* (*<u>DL</u>*<u>) approach.</u> For each drug on TCGA and HMF, we considered the architecture and the set of hyper-parameters with the lowest Mean Squared Error on GDSC given an initialization. We then randomly generated 50 independent initializations of the resulting networks and trained them using the GDSC data. Each of these trained networks was then employed to predict the TCGA or HMF response. The resulting prediction accuracies (area under the ROC) are plotted for the different drugs on the TCGA and HMF data. **A**: Deep Learning prediction accuracies for each drug on TCGA (green boxplots). For comparison, the results from TRANSACT are displayed (blue curve). **B**: Pearson correlation of the Mean Square Error of the predictor on GDSC to the Area under the ROC of the same predictor on TCGA. **C**: Boxplot of performances on each HMF drug for 50 different initializations. **D**: Pearson correlation on HMF between MSE (GDSC) and Area under the ROC (HMF).

**Supplementary Figure 12.**
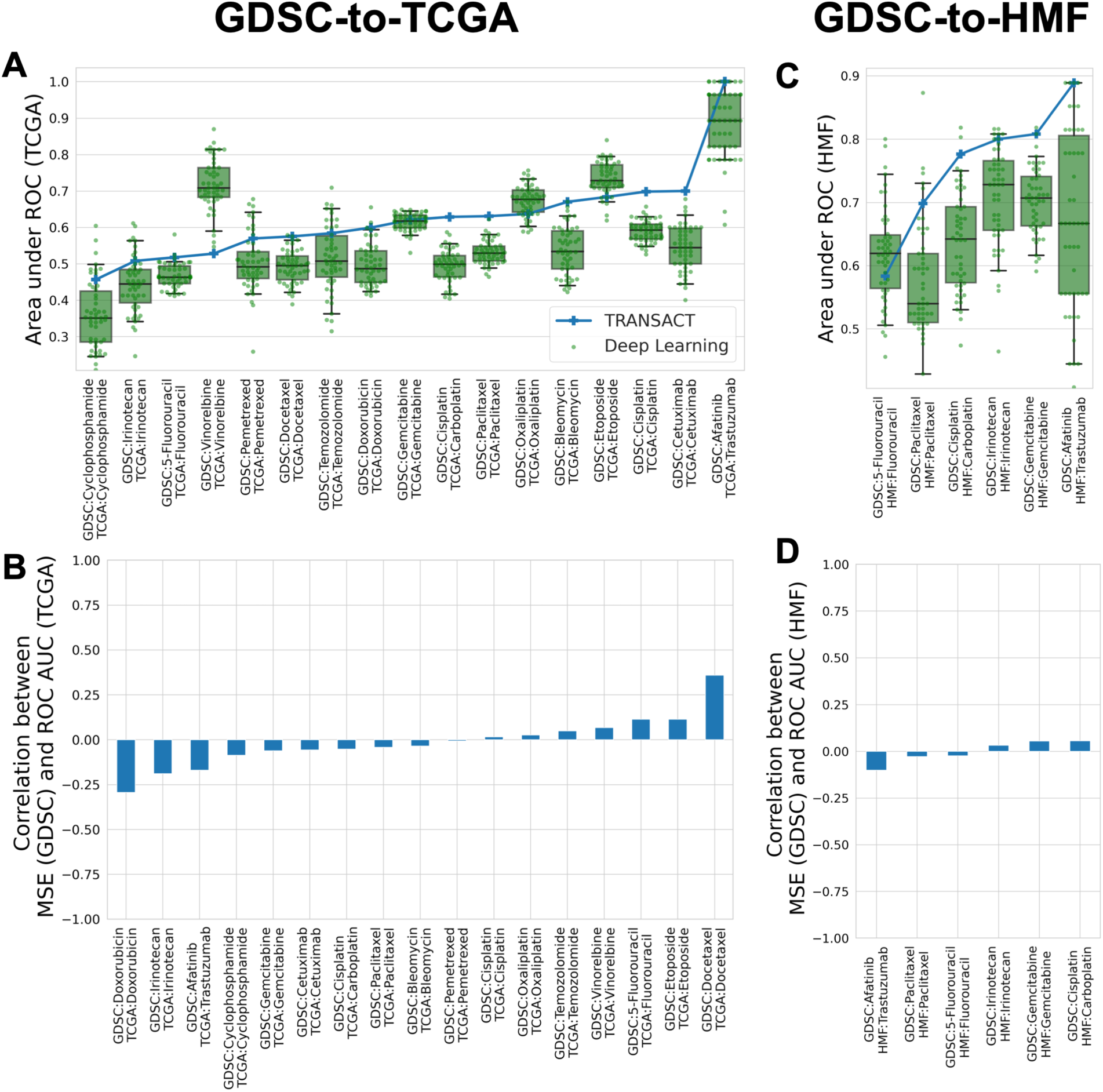
<u>Impact of initialization on results for the</u> *<u>ComBat</u>*<u>+</u>*<u>DL</u>* <u>approach.</u> For each drug on TCGA and HMF, we considered the architecture and the set of hyper-parameters with the lowest Mean Squared Error on GDSC given an initialization. We then randomly generated 50 independent initializations of the resulting networks and trained them using the GDSC data. Each of these trained networks was then employed to predict the TCGA or HMF response. The resulting predictions accuracies (area under the ROC) are plotted for the different drugs on the TCGA and HMF data. **A**: *ComBat+DL* prediction accuracies for each drug on TCGA (green boxplots). For comparison, the results from TRANSACT are displayed (blue curve). **B**: Pearson correlation of the Mean Square Error of the predictor on GDSC to the Area under the ROC of the same predictor on TCGA. **C**: Boxplot of performances on each HMF drug for 50 different initializations. **D**: Pearson correlation on HMF between MSE (GDSC) and Area under the ROC (HMF).

**Supplementary Figure 13.**
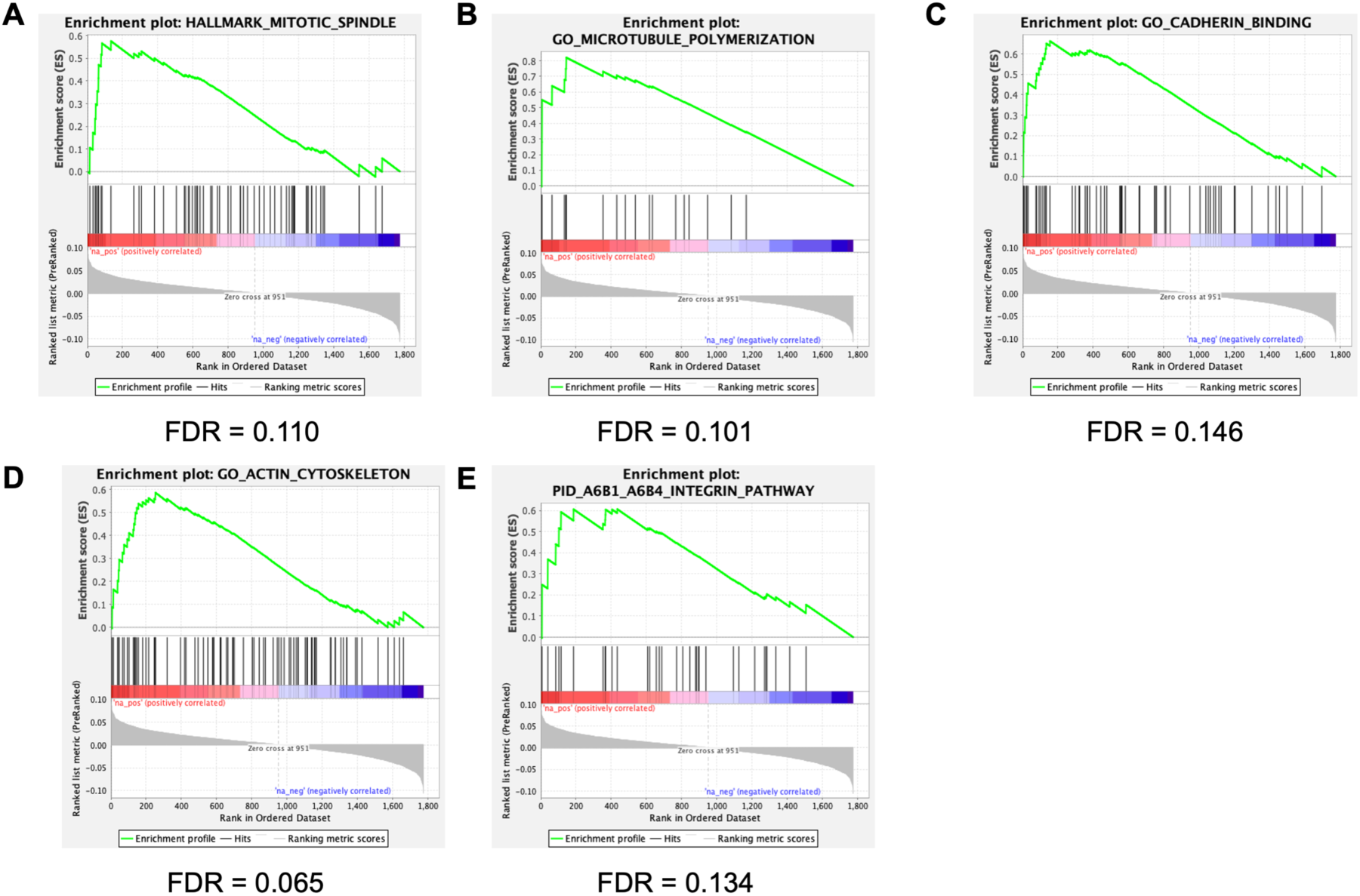
<u>Pathway enriched for resistant linear coefficients in GDSC-to-TCGA Gemcitabine drug response predictor</u>. Additional pathways significantly enriched in the linear part of the GDSC-to-TCGA predictor.

