## Supplementary material for "Predicting clinical drug response from model systems by non-linear subspace-based transfer learning": Algorithm Derivation

Soufiane Mourragui[1,2], Marco Loog[2,3], Daniel Vis[1], Kat Moore[1], Anna G Manjon[4], Mark A van de Wiel[5,6], Marcel JT Reinders[2,7,8] and Lodewyk FA Wessels[1,2,8]

[1]Division of Molecular Carcinogenesis, Oncode Institute, The Netherlands Cancer Institute, Amsterdam, The Netherlands.
[2]Department of EEMCS, Delft University of Technology, Delft, The Netherlands.
[3]Department of Computer Science, University of Copenhagen, Copenhagen, Denmark.
[4]Division of Cell Biology, Oncode Institute, The Netherlands Cancer Institute, Amsterdam, The Netherlands.
[5]Epidemiology and Biostatistics, Amsterdam University Medical Center, Amsterdam, The Netherlands.
[6]MRC Biostatistics Unit, Cambridge University, Cambridge, United Kingdom.
[7]Leiden Computational Biology Center, Leiden University Medical Center, Leiden, The Netherlands.

In this supplementary note, we present the algorithmic derivation of TRANSACT. Our approach works as follows:

1. We transform the original data (cell-view) and map it into a new space using a function $\varphi$. This mapping aims at representing the data in a more amenable way to standard linear analysis.

2. Once the whole dataset has been mapped, we find directions of importance in source and target datasets. Specifically, we reduce dimensionality, we align the low-rank directions, and interpolate between the two views to obtain single directions important in both datasets.

3. Finally, we project the mapped data in these directions. The obtained scores can then be used in any statistical model.

In the rest of this note, we prove the different steps leading to this extended algorithm. Although the note might seem technical, this all boils down to this overarching paradigm. To the reader who wishes to get directly to the main results, we highlighted the end products of our demonstration as Theorems (Theorems Supp 5.3, Supp 6.6 and Supp 8.5).

### Contents


**Supp 1 Notations and settings** **3**

**Supp 2 Kernel-mean centering** **3**

**Supp 3 Kernel PCA on source and target** **4**

**Supp 4 Variational definition of principal vectors** **5**

**Supp 5 Computation of Principal Vectors** **5**

**Supp 6 Interpolation scheme** **8**

**Supp 7 Gene set enrichment analysis of consensus features** **10**

**Supp 8 Equivalence with Geodesic Flow Kernel** **12**

**Supp 9 Difference with CCA on the genes** **13**

**Supp 10 Algorithm workflow** **14**

**Supp 11 Glossary** **15**

### Supp 1 Notations and settings

In our scenario, we have two datasets living in the same space – i.e. represented by the same $p$ features (genes, SNPs, methylation probes, ...):

- A source dataset $\mathcal{X}_s = \left\{x_1^s, x_2^s, .., x_{n_s}^s\right\} \subset \mathbb{R}^p$, with labels $\mathcal{Y}_s = \left\{y_1^s, ...y_{n_s}^s\right\}$.
- A target dataset $\mathcal{X}_t = \left\{x_1^t, x_2^t, .., x_{n_t}^t\right\} \subset \mathbb{R}^p$ usually unlabelled.

We represent the source (resp. target) data as a matrix $X_s \in \mathbb{R}^{n_s \times p}$ (resp. $X_t \in \mathbb{R}^{n_t \times p}$) with samples in the rows and features in the columns.
We consider a similarity function, or kernel, $K : \mathbb{R}^p \times \mathbb{R}^p \mapsto \mathbb{R}$ that we will assume for the sequel to be positive semi-definite. Using the theory of Reproducible Kernel Hilbert Space [1], $K$ is represented by the following dual formulation.

**Proposition Supp 1.1** (Reproducing Hilbert Space)**.** *There exists a unique functional Hilbert space* $(\mathcal{H}, \langle\cdot,\cdot\rangle_{\mathcal{H}})$*, with* $\mathcal{H} \subset \mathcal{F}(\mathbb{R}^p, \mathbb{R})$ *(functions from* $\mathbb{R}^p$ *to* $\mathbb{R}$*), and a mapping function* $\varphi : \mathbb{R}^p \mapsto \mathcal{H}$ *such that:*

$$\forall x, y \in \mathbb{R}^p, \quad K(x, y) = \langle \varphi(x), \varphi(y)\rangle_{\mathcal{H}}. \tag{Supp 1}$$

*The mapping* $\varphi$ *furthermore satisfies the Reproducing property:*

$$\forall f \in \mathcal{H}, \quad f : x \in \mathbb{R}^p \mapsto \langle \varphi(x), f\rangle_{\mathcal{H}}. \tag{Supp 2}$$

We refer to $d_s$ (resp. $d_t$) the number of low-rank components we reduced the source data (resp. target data) to. We set $d$ as the maximum number of principal vectors, $d = \min(d_s, d_t)$.
Superscript $s$ is used for source items and superscript $t$ for target items. $K(x, \cdot)$ , for $x \in \mathbb{R}^p$, is the function $y \in \mathbb{R}^p \mapsto K(x, y)$. We use the superscript $\cdot^T$ as the transposition operation.
Finally, we define the following kernel matrices:

**Definition Supp 1.2** (Kernel matrices)**.** *We define the following four matrices:*

- ***Source kernel matrix*** $K_s$ : $K_s = \left[K\left(x_i^s, x_j^s\right)\right]_{1\le i,j\le n_s} \in \mathbb{R}^{n_s \times n_s}$.
- ***Target kernel matrix*** $K_t$ : $K_t = \left[K\left(x_i^t, x_j^t\right)\right]_{1\le i,j\le n_t} \in \mathbb{R}^{n_t \times n_t}$.
- ***Source-target kernel matrix*** $K_{st}$ : $K_{st} = \left[K\left(x_i^s, x_j^t\right)\right]_{1\le i\le n_s, 1\le j\le n_t} \in \mathbb{R}^{n_s \times n_t}$.
- ***Target-source kernel matrix*** : $K_{ts}$ *as* $K_{ts} = K_{st}^T \in \mathbb{R}^{n_t \times n_s}$.

### Supp 2 Kernel-mean centering

We set out to work in the Hilbert space $\mathcal{H}$ after embedding the data with the mapping $\varphi$. Prior to any statistical processing, we first need to mean-center the data *in the kernel feature space* $\mathcal{H}$. For that purpose, we define two means, the *mean source embedding* $\mu^s$ and the *mean target embedding* $\mu^t$, as follows:

$$\begin{aligned} \mu^s &= \frac{1}{n_s}\sum_{i=1}^{n_s} \varphi\left(x_i^s\right) = \frac{1}{n_s}\sum_{i=1}^{n_s} K\left(x_i^s, \cdot\right) \\ \mu^t &= \frac{1}{n_t}\sum_{i=1}^{n_t} \varphi\left(x_i^t\right) = \frac{1}{n_s}\sum_{i=1}^{n_t} K\left(x_i^t, \cdot\right) \end{aligned} \tag{Supp 3}$$

Using the means computed in Equation (Supp 3), we define two sets of corrected embeddings as follows:

**Definition Supp 2.1** (Mean-centered embedding and kernel function). *The source centered kernel embedding* $\widetilde{\varphi}_s$ *is defined as:*

$$\forall x \in \mathbb{R}^p, \quad \widetilde{\varphi}_s(x) \;=\; \varphi(x) - \mu_s \;=\; K(x,\cdot) - \mu_s. \qquad \text{(Supp 4)}$$

*We then defined the source-centered kernel function* $\widetilde{K}_s$ *as:*

$$\forall x, y \in \mathbb{R}^p, \quad \widetilde{K}_s(x,y) \;=\; \langle \widetilde{\varphi}_s(x), \widetilde{\varphi}_s(y).\rangle \qquad \text{(Supp 5)}$$

*We define equivalently the target centered kernel embedding* $\widetilde{\varphi}_t$ *and corresponding target-centered kernel function* $\widetilde{K}_t$.

We use the mean-centered kernel functions defined in Definition Supp 2.1 to correct the kernel matrices from Definition Supp 1.2 and define the following four matrices.

**Definition Supp 2.2** (Centered Kernel matrices). *We define the following four matrices:*

- ***Source-centered kernel matrix*** $\widetilde{K}_s$ : $\widetilde{K}_s = \left[\widetilde{K}_s\left(x_i^s, x_j^s\right)\right]_{1\leq i,j\leq n_s} \in \mathbb{R}^{n_s\times n_s}$.
- ***Target-centered kernel matrix*** $\widetilde{K}_t$ : $\widetilde{K}_t = \left[\widetilde{K}_t\left(x_i^t, x_j^t\right)\right]_{1\leq i,j\leq n_t} \in \mathbb{R}^{n_t\times n_t}$.
- ***Source-target-centered kernel matrix*** $\widetilde{K}_{st}$ : $\widetilde{K}_{st} = \left[\langle \widetilde{\varphi}_s\left(x_i^s\right), \widetilde{\varphi}_t\left(x_j^t\right)\rangle\right]_{1\leq i\leq n_s, 1\leq j\leq n_t} \in \mathbb{R}^{n_s\times n_t}$.
- ***Target-source kernel matrix*** : $\widetilde{K}_{ts}$ *as* $\widetilde{K}_{ts} = \widetilde{K}_{st}^T \in \mathbb{R}^{n_t\times n_s}$.

To get a relation between matrices given in Definition Supp 2.2 and Definition Supp 1.2, we define the centering matrix of size $n$, denoted as $\mathbb{C}_n$:

**Definition Supp 2.3** (Centering matrix). *Let* $n \in \mathbb{N}_*$. *We define the centering matrix of size* $n$*, denoted* $C_n$ *as:*

$$C_n \;=\; I_n - \frac{1}{n}\mathbf{1}_n\mathbf{1}_n^T, \qquad \text{(Supp 6)}$$

*where* $I_n$ *is the identity matrix of size* $n$ *and* $\mathbf{1}_n$ *is the* $n$*-sized vector constituted solely of 1.*

**Proposition Supp 2.4** (Computation of centered kernel matrices). *We have the following equalities:*

$$\begin{aligned}\widetilde{K}_s &= C_{n_s}K_sC_{n_s},\\ \widetilde{K}_t &= C_{n_t}K_tC_{n_t},\\ \widetilde{K}_{st} &= C_{n_s}K_{st}C_{n_t}.\end{aligned} \qquad \text{(Supp 7)}$$

### Supp 3 Kernel PCA on source and target

We use Kernel PCA to compute directions of maximum variance in the embedded space [7], yielding kernel Principal Components, also called *non-linear principal components* (NLPCs) in the main text. These NLPCs for source and target are respectively defined as linear combinations of source and target samples' embeddings (after mean-centering) in the kernel feature space.

**Definition Supp 3.1** (Non-linear source and target principal components [7]). *Non-linear principal components for source* $\left(f_1^s, .., f_{d_s}^s\right)$ *and target* $\left(f_1^t, .., f_{d_t}^t\right)$ *are defined as linear combinations of source and target embedded samples respectively. Denoting as* $\alpha^s$ *the* $d_s$ *top eigenvectors of* $\widetilde{K}_s$ *and* $\alpha^t$ *the* $d_t$ *top eigenvectors of* $\widetilde{K}_t$*, we have the following equality:*

$$\begin{cases} f_q^s \;=\; \displaystyle\sum_{i=1}^{n_s} \alpha_{q,i}^s \widetilde{\varphi}_s\left(x_i^s\right) & \text{for } q \in \{1, .., d_s\}, \\ f_q^t \;=\; \displaystyle\sum_{i=1}^{n_t} \alpha_{q,i}^t \widetilde{\varphi}_t\left(x_i^t\right) & \text{for } q \in \{1, .., d_t\}, \end{cases} \qquad \text{(Supp 8)}$$

*These non-linear principal directions satisfy some orthogonality constraints on the kernel space $\mathcal{H}$:*

$$\forall x \in \{s,t\}\,, \quad \forall k,l \in \{1,..,d\}\,, \quad \langle f_k^x, f_l^x \rangle_{\mathcal{H}} = \delta_{k,l}, \tag{Supp 9}$$

*where $\delta$ is the equality indicator function. These constraints are equivalent to:*

$$\alpha^s \widetilde{K}_s {\alpha^s}^T = I_{d_s} \quad \textit{and} \quad \alpha^t \widetilde{K}_t {\alpha^t}^T = I_{d_t} \tag{Supp 10}$$

The two matrices $\alpha^s \in \mathbb{R}^{d_s \times n_s}$ and $\alpha^t \in \mathbb{R}^{d_t \times n_t}$ correspond to factors by samples matrices, but do not represent the projected score. Instead, they are equivalent to the feature loadings in linear PCA and correspond to a dual representation of the features in $\mathcal{H}$ that can not be explicitly computed due to the high-dimensions of $\mathcal{H}$. We refer to them as *sample importance loadings* to explicit the difference these have with projected scores.

### Supp 4 Variational definition of principal vectors

We define the first pair of principal vectors between source and target NLPCs as the two unitary vectors $s_1$ and $t_1$, with $s_1$ in source NLPCs span and $t_1$ in target NLPCs span, such that their similarity is maximized. This extends in $\mathcal{H}$ the principal vectors defined by Golub and Van Loan in [3] and are mathematically formalized using the following variational definition:

$$\begin{aligned} s_1, t_1 \quad = \quad & \underset{\substack{s \in \text{span}\left(f_1^s,..,f_{d_s}^s\right), \\ t \in \text{span}\left(f_1^t,..,f_{d_t}^t\right)}}{\text{argmax}} \quad \langle s,t \rangle_{\mathcal{H}} \\ & \text{s.t} \quad \langle s,s \rangle_{\mathcal{H}} = \langle t,t \rangle_{\mathcal{H}} = 1 \end{aligned} \quad . \tag{Supp 11}$$

We further define the principal vector by adding an orthogonality constraint, as in [3].

**Definition Supp 4.1** (Kernel Principal Vectors)**.** *We define the $d$ pairs of principal vectors $(s_1,t_1)\,,(s_2,t_2)\,,\,..,\,(s_d,t_d)$ as, for all $k \in \{1,..,d\}$ :*

$$\begin{aligned} s_k, t_k \quad = \quad & \underset{\substack{s \in \text{span}\left(f_1^s,..,f_{d_s}^s\right), \\ t \in \text{span}\left(f_1^t,..,f_{d_t}^t\right)}}{\text{argmax}} \quad \langle s,t \rangle_{\mathcal{H}} \\ & \textit{s.t} \quad \langle s,s \rangle_{\mathcal{H}} = \langle t,t \rangle_{\mathcal{H}} = 1, \\ & \textit{and } \forall l < k, s_l \perp s, t_l \perp t \end{aligned} \quad . \tag{Supp 12}$$

### Supp 5 Computation of Principal Vectors

The first step towards computing principal vectors is to compare the principal components defined in Definition Supp 3.1. We define for that purpose the cosine similarity matrix between source and target NLPCs and present a closed-form solution for computing it based on centered kernel matrices (Definition Supp 2.2) and NLPCs' coefficients (Definition Supp 3.1).
The cosine similarity matrix is a standard way to compare orthonormal basis of vectors and has already been used to compare linear principal components in subspace-based domain adaptation [2, 4, 5]. We here extend it to kernel-based non-linear dimensionality reduction.

**Definition Supp 5.1** (Cosine similarity matrix)**.** *We define the cosine similarity matrix $\mathbf{M}^K$ between source and target kernel principal components as:*

$$\mathbf{M}^K \quad = \quad \left[\langle f_k^s, f_l^t \rangle_{\mathcal{H}}\right]_{1 \le k \le d_s, 1 \le l \le d_t} \quad \in \mathbb{R}^{d_s \times d_t}. \tag{Supp 13}$$

**Proposition Supp 5.2** (Computation of cosine similarity matrix). *$\mathbf{M}^K$ can be computed using the matrices $\alpha^s$, $\alpha^t$ and $K_{ST}$ as:*

$$\begin{aligned}\mathbf{M}^K &= \alpha^s \widetilde{K}_{st} {\alpha^t}^T \\ &= \alpha^s C_{n_s} K_{st} C_{n_t} {\alpha^t}^T.\end{aligned} \qquad \text{(Supp 14)}$$

*Proof.* Let $1 \leq k, l \leq d$, then using Equation Supp 8,

$$\langle f_k^s, f_l^t \rangle = \sum_{i=1}^{n_s} \sum_{j=1}^{n_t} \alpha_{k,i}^s \alpha_{l,j}^t \langle \widetilde{\varphi}_s(x_i^s), \widetilde{\varphi}_t(x_j^t) \rangle = {\alpha_{k,:}^s}^T \widetilde{K}_{st} \alpha_{l,:}^t \qquad \text{(Supp 15)}$$

which put together as a matrix gives the wanted result. ■

Similarly to the linear setting, we use this cosine similarity matrix to NLPC by means of SVD of $\mathbf{M}^K$.

**Theorem Supp 5.3** (SVD computation of Principal Vectors). *Let $\beta^s \in \mathbb{R}^{d_s \times d}$ (resp. $\beta^t \in \mathbb{R}^{d_t \times d}$) be the first $d$ left (resp. right) singular vectors of $\mathbf{M}^K$, i.e. $\mathbf{M}^K \approx \beta^s \Sigma {\beta^t}^T$. Then, for all $1 \leq q \leq d$:*

$$s_q = \sum_{k=1}^{d_s} \sum_{i=1}^{n_s} \beta_{k,q}^s \alpha_{k,i}^s \widetilde{\varphi}_s(x_i^s) \quad \textit{and} \quad t_q = \sum_{l=1}^{d_t} \sum_{j=1}^{n_t} \beta_{l,q}^t \alpha_{l,j}^t \widetilde{\varphi}_t(x_j^t) \qquad \text{(Supp 16)}$$

*Proof.* Let $s_1, .., s_d \in \text{span}\left(f_1^s, .., f_{d_s}^s\right)$ and $t_1, .., t_d \in \text{span}\left(f_1^t, .., f_{d_t}^t\right)$ with norm 1, there exists $\beta^s, \in \mathbb{R}^{d_s \times d}$ and $\beta^t \in \mathbb{R}^{d_t \times d}$ such that, for all $q \in \{1, .., d\}$,

$$s_q = \sum_{k=1}^{d_s} \beta_{k,q}^s f_k^s = \sum_{i=1}^{n_s} \sum_{k=1}^{d_s} \alpha_{k,i}^s \beta_{k,q}^s \widetilde{\varphi}_s(x_i^s) \quad \text{and} \quad t_q = \sum_{l=1}^{d_t} \beta_{l,q}^t f_l^t = \sum_{j=1}^{n_t} \sum_{l=1}^{d_t} \alpha_{l,j}^t \beta_{l,q}^t \widetilde{\varphi}_t(x_j^t). \qquad \text{(Supp 17)}$$

The orthogonality constraint $\langle s_k, s_l \rangle_{\mathcal{H}} = \langle t_k, t_l \rangle_{\mathcal{H}} = \delta_{k,l}$, for $1 \leq k, l \leq d$ coupled with the orthogonality constrains from Equation (Supp 9) is then equivalent to ${\beta^s}^T \beta^s = {\beta^t}^T \beta^t = I_d$. Computing inner product between source and target PV therefore yields

$$[\langle s_k, t_l \rangle]_{1 \leq k,l \leq d} = {\beta^s}^T \alpha^s \widetilde{K}_{st} {\alpha^t}^T {\beta^t}^T = {\beta^s}^T \mathbf{M}^K \beta^t. \qquad \text{(Supp 18)}$$

Therefore, the maximization problem from Equation (Supp 11) is equivalent to the following:

$$\begin{aligned}\max_{\substack{\beta^s \in \mathbb{R}^{d_s \times d}, \\ \beta^t \in \mathbb{R}^{d_t \times d}}} \quad & {\beta^s}^T \mathbf{M}^K \beta^t \\ \text{s.t.} \quad & {\beta^s}^T \beta^s = {\beta^t}^T \beta^t = I_d\end{aligned}, \qquad \text{(Supp 19)}$$

which unique solutions are the left and right orthogonal vectors of $\mathbf{M}^K$, obtained by SVD. ■

In order to work at the sample-level for each principal vector, we define the PV sample-importance loadings as follows.

**Definition Supp 5.4** (Principal Vector sample importance loadings). *We define the source (resp. target) sample importance loadings $\rho^s \in \mathbb{R}^{d \times n_s}$ (resp. $\rho^t \in \mathbb{R}^{d \times n_t}$) as:*

$$\rho^s = {\beta^s}^T \alpha^s \quad \textit{and} \quad \rho^t = {\beta^t}^T \alpha^t. \qquad \text{(Supp 20)}$$

These PV importance loadings are related to the source and target PVs as follow:

**Proposition Supp 5.5.** *Source and target principal vectors have the equivalent following definition:*

$$\forall q \in \{1, ..d\}, \quad \begin{cases} s_q &= \sum_{i=1}^{n_s} \rho^s_{q,i} \widetilde{\varphi}_s\left(x^s_i\right), \\ t_q &= \sum_{i=1}^{n_t} \rho^t_{q,i} \widetilde{\varphi}_t\left(x^t_i\right). \end{cases} \tag{Supp 21}$$

We finally defined the similarity between the principal vectors as cosines of angles referred to as *principal angles.*

**Definition Supp 5.6** (Principal Angles)**.** *Let $1 \leq q \leq d$. We define the q-th principal angle as the unique $\theta_q \in \left[0, \frac{\pi}{2}\right]$ that satisfies:*

$$\cos\theta_q = \langle s_q, t_q \rangle_{\mathcal{H}}. \tag{Supp 22}$$

**Proposition Supp 5.7** (SVD computation of Principal Angles)**.** *Let $\Sigma$ be the diagonal matrix obtained by SVD of $\mathbf{M}^K$ (as in Proposition Supp 5.2), then:*

$$\forall q \in \{1, .., d\}, \quad \cos\theta_q = \Sigma_{q,q}. \tag{Supp 23}$$

*Proof.*

$$\cos\theta_q = \langle s_q, t_q \rangle_{\mathcal{H}} = {\beta^s_{:,q}}^T \mathbf{M}^K \beta^t_{:,q} = \Sigma_{q,q}, \tag{Supp 24}$$

by definition of the SVD. ■

We showed how to compute the PVs as functions in $\mathcal{H}$ and gave a closed-form solution for the evaluation in $\mathbb{R}^p$. We finally show that the evaluation of PVs correspond to a projection of the embedded vector, keeping the same intuition than in linear setting.

**Proposition Supp 5.8** (Evaluation of principal vectors)**.** *Let $x \in \mathbb{R}^p$. For $q \in \{1, ..d\}$, the evaluation of source and target principal vectors $s_q$ and $t_q$ is equivalent to the projection of the embedding of $x$ on these vectors:*

$$s_q(x) = \langle s_q, \varphi(x) \rangle_{\mathcal{H}} \quad \text{and} \quad t_q(x) = \langle t_q, \varphi(x) \rangle_{\mathcal{H}} \tag{Supp 25}$$

*Proof.* Combining Equations (Supp 3), (Supp 4) and (Supp 21), source PV are sum of elements of $\mathcal{H}$:

$$s_q = \sum_{i=1}^{n_s} \rho^s_{q,i} \widetilde{\varphi_s}\left(x^s_i\right) \quad \text{with,} \quad \forall i \in \{1, .., n_s\}, \ \widetilde{\varphi_s}\left(x^s_i\right) \in \mathcal{H}. \tag{Supp 26}$$

Therefore $s_q \in \mathcal{H}$ since $\mathcal{H}$ is an Hilbert space. Using the reproducing property of the RKHS and the definition of $\varphi$ (Equation (Supp 1), we obtain

$$\forall x \in \mathbb{R}^p, \quad s_q(x) = \langle s_q, \varphi(x) \rangle_{\mathcal{H}}. \tag{Supp 27}$$

Following the same idea, we obtain the equivalent equality for target PVs. ■

#### Supp 6 Interpolation scheme

The Principal Vectors are pairs of vectors (one form source, one from target) that are geometrically similar. We select only the pairs above a certain threshold of similarity in order to restrict to directions shared by the two signals. Therefore, within each pair, source and target vectors show an important correlation and using the two into a predictive model would not be optimal. We therefore set out to construct a single vector out of each pair by interpolating between the two vectors. This interpolation is the geodesic flow between PVs and is defined as follows.

**Definition Supp 6.1** (Angular interpolation function)**.** *Let $q \in \{1, .., d\}$, we define the angular interpolation functions $\Gamma_q$ and $\xi_q$ between the $q^{th}$ pair of principal vector as:*

$$\forall \tau \in [0,1], \quad \Gamma_q(\tau) = \frac{\sin((1-\tau)\theta_q)}{\sin\theta_q} \quad \text{and} \quad \xi_q(\tau) = \frac{\sin \tau\theta_q}{\sin\theta_q}. \tag{Supp 28}$$

**Definition Supp 6.2** (Geodesic flow between principal vectors)**.** *Let $q \in \{1, .., d\}$, we define the interpolation $\phi_q$ between the $q^{th}$ pair of principal vector as:*

$$\forall \tau \in [0,1], \quad \phi_q(\tau) \ = \ \Gamma_q(\tau)\, s_q \ + \ \xi_q(\tau)\, t_q. \tag{Supp 29}$$

*Since $\mathcal{H}$ is a Hilbert space, $\phi_q \in \mathcal{H}$.*

**Proposition Supp 6.3** (Estimation using PV sample importance loadings)**.** *Let $q \in \{1, .., d\}$ and $\phi_q$ be the geodesic between the $q^{th}$ pair of principal vectors. The geodesic defined in Equation (Supp 29) has the following equivalent formulation:*

$$\forall \tau \in [0,1], \quad \phi_q(\tau) \ = \ \Gamma_q(\tau) \sum_{i=1}^{n_s} \rho^s_{q,i} \widetilde{\varphi}_s(x^s_i) \ + \ \xi_q(\tau) \sum_{j=1}^{n_t} \rho^t_{q,j} \widetilde{\varphi}_t\left(x^t_j\right). \tag{Supp 30}$$

*Proof.* Combining the definition of the geodesic from Definition Supp 6.2 with the equivalent principal vector formulation of Proposition Supp 5.5 yields the result. ■

The formulation of the geodesic from Proposition Supp 6.3 can easily be written down as a matrix product (for computation purposes) for each sample. We define the matrix angular interpolation function as follow.

**Definition Supp 6.4** (Matrix angular interpolation function)**.** *We define the matrix angular interpolation functions $\mathbf{\Gamma}$ and $\mathbf{\Xi}$*

$$\forall \tau \in [0,1]^d, \quad \mathbf{\Gamma}(\tau) = \text{diag}\left[\Gamma_q(\tau_q)\right]_{1\leq q\leq d} \quad \text{and} \quad \mathbf{\Xi}(\tau) = \text{diag}\left[\xi(\tau_q)\right]_{1\leq q\leq d}. \tag{Supp 31}$$

**Proposition Supp 6.5** (Matrix estimation of principal vectors)**.** *Let's denote by $s$ (resp. $t$) the vectors of $d$ source (resp. target) principal vectors ordered by similarity. We define $\mathcal{S}^s$ and $\mathcal{S}^t$ as the matrices that contain the source principal vectors values evaluated on source and target data respectively:*

$$\mathcal{S}^s = \left[s\left(x^s_1\right)^T, .., s\left(x^s_{n_s}\right)^T\right]^T \in \mathbb{R}^{n_s \times d}, \tag{Supp 32}$$

$$\mathcal{S}^t = \left[s\left(x^t_1\right)^T, .., s\left(x^t_{n_t}\right)^T\right]^T \in \mathbb{R}^{n_t \times d}. \tag{Supp 33}$$

*We define similarly $\mathcal{T}^s \in \mathbb{R}^{n_s \times d}$ as the matrix that contains the target principal vectors evaluated on the source data – and $\mathcal{T}^t \in \mathbb{R}^{n_t \times d}$ as the matrix that contains the target principal vectors evaluated on the target data. These matrices can be computed as follows:*

$$\begin{cases} \mathcal{S}^s &= K^s C_{n_s} {\rho^s}^T, \\ \mathcal{S}^t &= K^{st} C_{n_s} {\rho^s}^T, \end{cases} \quad \textit{and} \quad \begin{cases} \mathcal{T}^t &= K^{ts} C_{n_t} {\rho^t}^T. \\ \mathcal{T}^t &= K^t C_{n_t} {\rho^t}^T. \end{cases} \tag{Supp 34}$$

*Proof.* Using the definition of principal vectors with $\rho$ coefficients from Equation (Supp 21), we get, for $l \in \{1, .., n_s\}$ and $q \in \{1, .., d\}$:

$$\begin{aligned} s_q\left(x_l^s\right) &= \sum_{i=1}^{n_s} \rho_{q,i}^s \left[ K\left(x_i^s, x_l^s\right) - \frac{1}{n_s} \sum_{j=1}^{n_s} \left(x_j^s, x_l^s\right) \right] \\ &= \sum_{i=1}^{n_s} \left(\rho_{q,:}^s\right)_i \left[ K_{i,l}^s - \frac{1}{n_s} \left(\mathbf{1}_{n_s} \mathbf{1}_{n_s}^T K^s\right)_{i,l} \cdot \right] \end{aligned} \tag{Supp 35}$$

Using the centering matrix defined in Definition Supp 2.3, we get:

$$s_k\left(x_l^s\right) = \left[\rho^s C_{n_s} K^s\right]_{k,l}, \tag{Supp 36}$$

and therefore $\mathcal{S}^s = \left(\rho^s C_{n_s} K^s\right)^T$. The other equalities follow from the same proof.

■

Let's finally define the geodesic matrix between source and target at interpolation time $\tau \in [0,1]$ as the estimation of both source and target on the geodesic in the kernel feature space.

**Theorem Supp 6.6.** *We define as $\mathbf{F}^{st}(\tau)$ for as the matrix of geodesic values evaluated at interpolation time $\tau \in [0,1]^d$, i.e.:*

$$\mathbf{F}^{st}(\tau) = \begin{bmatrix} \phi(\tau)\left(x_1^s\right) \\ \cdots \\ \phi(\tau)\left(x_{n_s}^s\right) \\ \phi(\tau)\left(x_1^t\right) \\ \cdots \\ \phi(\tau)\left(x_{n_t}^s\right) \end{bmatrix} \in \mathbb{R}^{(n_s+n_t)\times d}. \tag{Supp 37}$$

*. Then $\mathbf{F}^{st}(\tau)$ can be computed as follow:*

$$\mathbf{F}^{st}(\tau) = \begin{bmatrix} \mathcal{S}^s & \mathcal{T}^s \\ \mathcal{S}^t & \mathcal{T}^t \end{bmatrix} \begin{bmatrix} \mathbf{\Gamma}(\tau) \\ \mathbf{\Xi}(\tau) \end{bmatrix}. \tag{Supp 38}$$

*This formulation is equivalent to:*

$$\mathbf{F}^{st}(\tau) = \begin{bmatrix} K^s & K^{st} \\ K^{ts} & K^t \end{bmatrix} \begin{bmatrix} C_{n_s} & 0_{n_s \times n_t} \\ 0_{n_t \times n_s} & C_{n_t} \end{bmatrix} \begin{bmatrix} {\rho^s}^T & 0_{n_s \times d} \\ 0_{n_t \times d} & {\rho^t}^T \end{bmatrix} \begin{bmatrix} \mathbf{\Gamma}(\tau) \\ \mathbf{\Xi}(\tau) \end{bmatrix}. \tag{Supp 39}$$

*Proof.* Direct by combining Definition Supp 6.4 and Proposition Supp 6.5. ■

In order to get zero-centered projected source and target samples, we can use two solutions. On one hand, we can perform a consensus-feature-level mean-centering independently on source and target after projection. Equivalently, we can also left-multiply by centering matrix the projected matrix $\mathbf{F}^{st}(\tau)$.
We finally show that the evaluation of the consensus features functions is equivalent to the projection of embedding in the feature space $\mathcal{H}$.

**Proposition Supp 6.7.** *Let $x \in \mathbb{R}^p$, $q \in \{1,..d\}$ and $\tau_q \in [0,1]$, then:*

$$\phi_q(\tau_q)(x) = \langle \phi_q(\tau_q), \varphi(x) \rangle_{\mathcal{H}}. \qquad \text{(Supp 40)}$$

*Proof.* Using Proposition Supp 5.8,

$$\begin{aligned} \phi_q(\tau_q)(x) &= \Gamma_q(\tau_q) s_q(x) + \xi_q(\tau_q) t_q(x) \\ &= \langle \Gamma_q(\tau_q) s_q + \xi_q(\tau_q) t_q, \varphi(x) \rangle_{\mathcal{H}} \\ &= \langle \phi_q(\tau_q), \varphi(x) \rangle_{\mathcal{H}}. \end{aligned} \qquad \text{(Supp 41)}$$

■

### Supp 7 Gene set enrichment analysis of consensus features

In order to gain insight into the making of consensus features, we use a Taylor expansion of the Gaussian kernel [8]. The Gaussian kernel can be expressed as outer-product of the following basis functions.

**Definition Supp 7.1.** *Let $i \leq 0$ be an integer. We define as $e_i : \mathbb{R} \mapsto \mathbb{R}$ the basis function defined as:*

$$\forall x \in \mathbb{R}, \quad e_i(x) = \sqrt{\frac{2\gamma^i}{i!}} x^i \exp\left(-\gamma x^2\right). \qquad \text{(Supp 42)}$$

**Proposition Supp 7.2** (Countable orthonormal basis of $\mathcal{H}$ [8])**.** *Let's define for $(i_1,..,i_p) \in \mathbb{N}^p$ the following function*

$$e_{(i_1,..,i_p)} = x \in \mathbb{R}^p \mapsto e_{i_1}(x_1) \times e_{i_2}(x_2) \times \cdot \cdot \times e_{i_p}(x_p). \qquad \text{(Supp 43)}$$

*Then, $\left(e_{(i_1,..,i_p)}\right)_{(i_1,..,i_p) \in \mathbb{N}^p}$ is an orthonormal basis of $\mathcal{H}$, and for $x, y \in \mathbb{R}^p$,*

$$\begin{aligned} \exp\left(-\gamma ||x-y||^2\right) &= \sum_{i_1,..,i_p \in \mathbb{N}^p} e_{(i_1,..,i_p)}(x) \, e_{(i_1,..,i_p)}(y). \\ &= \widehat{\varphi}(x)^T \widehat{\varphi}(y), \end{aligned} \qquad \text{(Supp 44)}$$

*with $\widehat{\varphi} : x \mapsto \left(e_{(i_1,..,i_p)}(x)\right)_{(i_1,..,i_p) \in \mathbb{N}^p}$.*

Let's consider this approximation map $\widehat{\varphi}$. We extract three different features of interest for our analysis: the offset (sum of indices is 0), the linear terms (sum of indices is 1) and the interaction terms (sum of indices is 2). We define them as follows:

**Definition Supp 7.3** (Offset, linear and interaction terms)**.**
*We define the **offset feature** $e_O$ as $e_{0_{\mathbb{N}^p}}$, i.e. when all indices are 0.*
*For each gene (feature $k \in \{1,..,p\}$), we define the $k^{th}$ **linear feature** $e_k$ as $e_{\delta_k}$ where $\delta_k$ is the vector of zeros with a single 1 on $k^{th}$ position.*
*For each combination of genes (feature $k, l \in \{1,..,p\}$), we define the $(k,l)^{th}$ **interaction feature** $e_{k,l}$ as $e_{\delta_{k,l}}$ where $\delta_{k,l}$ is the vector of zero with one 1 on $k^{th}$ and $l^{th}$ position only if $k \neq l$, and 2 on $k^{th}$ position if $k = l$.*

**Definition Supp 7.4** (Offset, linear and interaction terms for consensus features)**.**
*We define the **offset contribution** to consensus feature $q$ as $\mathcal{O}_q = \langle e_O, \phi_q(\tau_q^*) \rangle$.*
*For $k \in \{1,..,p\}$, we define the $k^{th}$ **linear contribution** to consensus feature $q$ as $\mathcal{L}_{q,k} = \langle e_k, \phi_q(\tau_q^*) \rangle$.*
*For $k, l \in \{1,..,p\}$), we define the $(k,l)^{th}$ **interaction contribution** to consensus feature $q$ as $\mathcal{I}_{q,k,l} = \langle e_{k,l}, \phi_q(\tau_q^*) \rangle$.*

We now compute the contribution of each of these features to the consensus features. We first rewrite the different contributions to the consensus features for readability.

**Definition Supp 7.5.** *For $q \in \{1,..,d\}$, we define $\sigma_q^s = \Gamma_q\left(\tau_q^*\right)\rho_q^s$ and $\sigma_q^t = \xi_q\left(\tau_q^*\right)\rho_q^t$.*

We finally define the source and target mean centered features.

**Definition Supp 7.6.** *We define the source (resp. target) mean-centered offset feature for the $q^{th}$ consensus feature $\widetilde{e}_O^s$ (resp. $\widetilde{e}_O^t$) as:*

$$\widetilde{e}_O^s = e_O - \frac{1}{n_s}\sum_{i=1}^{n_s} e_O\left(x_i^s\right) \quad \text{and} \quad \widetilde{e}_O^t = e_O - \frac{1}{n_t}\sum_{i=1}^{n_t} e_O\left(x_i^t\right). \tag{Supp 45}$$

*For $k \in \{1,..,p\}$, we define the source (resp. target) mean-centered linear feature for the $q^{th}$ consensus feature $\widetilde{e}_k^s$ (resp. $\widetilde{e}_k^t$) as:*

$$\widetilde{e}_k^s = e_k - \frac{1}{n_s}\sum_{i=1}^{n_s} e_k\left(x_i^s\right) \quad \text{and} \quad \widetilde{e}_k^t = e_k - \frac{1}{n_t}\sum_{i=1}^{n_t} e_k\left(x_i^t\right). \tag{Supp 46}$$

*For $k,l \in \{1,..,p\}$, we define the source (resp. target) mean-centered linear feature for the $q^{th}$ consensus feature $\widetilde{e}_{k,l}^s$ (resp. $\widetilde{e}_{k,l}^t$) as:*

$$\widetilde{e}_{k,l}^s = e_{k,l} - \frac{1}{n_s}\sum_{i=1}^{n_s} e_{k,l}\left(x_i^s\right) \quad \text{and} \quad \widetilde{e}_{k,l}^t = e_{k,l} - \frac{1}{n_t}\sum_{i=1}^{n_t} e_{k,l}\left(x_i^t\right). \tag{Supp 47}$$

**Proposition Supp 7.7.** *The differents contribution $\mathcal{O}_q$ $\mathcal{L}_{q,i}$ and $\mathcal{I}_{q,i,j}$ for the $q^{th}$ consensus feature can be computed as follow:*

$$\mathcal{O}_q = \sum_{i=1}^{n_s}\sigma_{q,i}^s\widetilde{e}_O^s\left(x_i^s\right) + \sum_{i=1}^{n_t}\sigma_{q,i}^t\widetilde{e}_O^t\left(x_i^t\right), \tag{Supp 48}$$

$$\mathcal{L}_{q,k} = \sum_{i=1}^{n_s}\sigma_{q,i}^s\widetilde{e}_{q,k}^s\left(x_i^s\right) + \sum_{i=1}^{n_t}\sigma_{q,i}^t\widetilde{e}_{q,k}^t\left(x_i^t\right), \tag{Supp 49}$$

$$\mathcal{I}_{q,k,l} = \sum_{i=1}^{n_s}\sigma_{q,i}^s\widetilde{e}_{q,k,l}^s\left(x_i^s\right) + \sum_{i=1}^{n_t}\sigma_{q,i}^t\widetilde{e}_{q,k,l}^t\left(x_i^t\right). \tag{Supp 50}$$

*Proof.* Combining the expression of consensus features as mean-centered source and target embedding from Supp 6.3, Definition Supp 7.5 and Definitions Supp 7.4 and Supp 7.6 gives the wanted results. ■

**Definition Supp 7.8.** *For the $q^{th}$ consensus feature, we define the* ***offset proportion*** *as $O_q = \mathcal{O}_q^2$, the* ***linear contribution*** *as $L_q = \sum_{k=1}^{p}\mathcal{L}_{q,k}^2$ and the* ***interaction contribution*** *as $I_q = \sum_{1\leq k\leq l\leq p}\mathcal{I}_{q,k,l}^2$.*
*Finally, we define the* ***higher-order contribution*** *as $R_q = 1 - O_q - L_q - I_q$.*

We now restrict to one gene set to measure the effect of this gene set on interactions and linear effects.
We here restricted to the Gaussian kernel. However, our results would easily be extended to any kernel, provided the feature space $\mathcal{H}$ has a known orthonormal basis.

### Supp 8 Equivalence with Geodesic Flow Kernel

In this section we showed the equivalence with the previously published linear version of the algorithm, the so-called PRECISE model [6]. We recall the main steps of the algorithm.

**Definition Supp 8.1** (Linear Principal Vectors). *Let $P_s \in \mathbb{R}^{d_s \times p}$ and $P_t \in \mathbb{R}^{d_t \times p}$ be two families of orthonormal vectors, i.e. $P_s P_s^T = I_{d_s}$ and $P_t P_t^T = I_{d_t}$. We define the cosine similarity matrix $\mathbf{M}$ as:*

$$\mathbf{M} \;=\; P_s P_t^T. \tag{Supp 51}$$

*Let $d \leq \min(d_s, d_t)$ and let $U\Sigma^L V^T$ be the $d-$ rank SVD approximation of $M$. We define the $d$ source (resp. target) principal vectors as the matrix $\mathbf{Q}_s \in \mathbb{R}^{d \times p}$ (resp. $\mathbf{Q}_t \in \mathbb{R}^{d \times p}$) as:*

$$\mathbf{Q}_s \;=\; U^T P_s \quad \textit{and} \quad \mathbf{Q}_t \;=\; V^T P_t. \tag{Supp 52}$$

*Samples can be projected on these four matrices ($P_s, P_t, \mathbf{Q}_s$ and $\mathbf{Q}_t$) by inner-product, i.e. canonical projection operator in Euclidean space.*

$P_s$ and $P_t$ are here defined generally as two families of orthonormal vectors. In particular, we consider for the rest that they are the results of PCA on respectively the source and the target covariance matrices $X_s^T C_{n_s} C_{n_s}^T X_s$ and $X_t^T C_{n_t} C_{n_t}^T X_t$. Using the linear PVs from Definition Supp 8.1, we define a linear interpolation scheme as follows.

**Definition Supp 8.2** (Linear Interpolation). *Using notations from Definition Supp 8.1, we define the linear principal angles as:*

$$\forall q \in \{1, .., d\}, \quad \cos\theta_q^L = \Sigma_{q,q}^L. \tag{Supp 53}$$

*For the PV pair $q \in \{1, .., d\}$, we define the interpolation function $\phi_q^L$ as follows:*

$$\phi_q : \; \tau_q \in [0,1] \mapsto \frac{\sin(1-\tau_q)\theta_q}{\sin\theta_q}(Q_s)_q \;+\; \frac{\sin\tau_q\theta_q}{\sin\theta_q}(Q_t)_q \tag{Supp 54}$$

Before stating the main result, we need the following well-known lemma.

**Lemma Supp 8.3** (Equivalence of spectrum). *Let $X \in \mathbb{R}^{n \times p}$. We denote by $S^+ = \left\{\left(\lambda_1^+, v_1^+\right), .., \left(\lambda_{d^+}^+, v_{d^+}^+\right)\right\}$ the non-singular spectrum of $XX^T$ and $S^- = \left\{\left(\lambda_1^-, v_1^-\right), .., \left(\lambda_{d^-}^-, v_{d^-}^-\right)\right\}$ the non-singular spectrum of $X^T X$, i.e. $\lambda_1 \geq \lambda_2 \geq .. \geq \lambda_d > 0$ in both spectrum. Then $d^+ = d^-$ and*

$$\forall i \in \left\{1, .., d^+\right\}, \quad \lambda_i^+ \;=\; \lambda_i^-, \quad v_i^+ = X v_i^- \quad \textit{and} \quad v_i^- = X^T v_i^+ \tag{Supp 55}$$

We consider two scenario using the same source and target datasets: linear PRECISE, and our kernelized approach with a linear kernel. We consider all other parameters set to the same values.

**Proposition Supp 8.4** (Equality of cosine similarity matrixes). *Let $\mathbf{M}$ and $\mathbf{M}^K$ be the cosine similarity matrices obtained respectively using linear PRECISE (Definition Supp 8.1) and the kernelized version with a linear kernel (Definition Supp 5.1), all hyperparameters equal. Then $\mathbf{M} = \mathbf{M}^\mathbf{K}$.*

*Proof.* We here use notations from Definitions Supp 5.1 and Supp 3.1. We define $\widetilde{X_s} = C_{n_s} X_s$ and $\widetilde{X_t} = C_{n_t} X_t$. We also use $\widetilde{K_s} = \widetilde{X_s}\widetilde{X_s}^T$ and $\widetilde{K_t} = \widetilde{X_t}\widetilde{X_t}^T$.

By definition of PCA, $P_s$ contains the top $d_s$ eigenvectors of the matrix $\widetilde{X_s}^T \widetilde{X_s}$, while $\alpha^s \widetilde{K_s}^{\frac{1}{2}}$ contains the top $d_s$ eigenvectors of $\widetilde{X_s}\widetilde{X_s}^T$. Using the result from Supp 8.3, we have $P_s = \alpha^s \widetilde{K_s}^{\frac{1}{2}} \widetilde{X_s}$. Similarly, we obtain on the target $P_t = \alpha^t \widetilde{K_t}^{\frac{1}{2}} \widetilde{X_t}$. Using Proposition Supp 5.2,

$$\mathbf{M} \;=\; \alpha^s \widetilde{K_s}^{\frac{1}{2}} \widetilde{X_s}\widetilde{X_t}^T \widetilde{K_t}^{\frac{1}{2}} \alpha^{t^T} \;=\; \alpha^s \widetilde{X_s}\widetilde{X_t}^T \alpha^{t^T} \;=\; \mathbf{M}^K, \tag{Supp 56}$$

using the fact that $\alpha^s \widetilde{K_s}^{\frac{1}{2}}$ and $\alpha^t \widetilde{K_t}^{\frac{1}{2}}$ is an eigenvector of $\widetilde{K_s}$ and $\widetilde{K_t}$ respectively. ■

From Proposition Supp 8.4 follows directly this equivalence.

**Theorem Supp 8.5.** *With all hyperparameters equal, PRECISE and the kernelized version with a linear kernel are equivalent.*

Theorem Supp 8.5 shows that all results obtained in linear case [6] hold for TRANSACT with a linear similarity function, and in particular the correspondence with the Geodesic Flow Kernel.

### Supp 9 Difference with CCA on the genes

Another data-strategy used in single-cell data analysis consists in using the gene-level correspondence to perform a Canonical Correlation Analysis (CCA) on the genes. Using the same notations as in Section Supp 1, this approach boils down to solve the following maximization procedure:

$$s_1, t_1 = \underset{\substack{s \in \mathbb{R}^{n_s}, s^T s = 1, \\ t \in \mathbb{R}^{n_t}, t^T t = 1}}{\text{argmax}} \; s^T X_s X_t^T t \qquad \text{(Supp 57)}$$

and the subsequent directions defined orthogonally to these directions. This procedure find directions of maximum covariance at the gene-level between source and target. It will find two combinations of samples (one for source, and one for target) that show the maximum covariance among genes. It differs markedly from our methods on several aspects. First, from a computational standpoint, the SVD-equivalent definition of PVs (Theorem Supp 5.3) consists in breaking down a relatively small matrix ($d_s \times d_t$) and not a sample-sample similarity matrix. Second, by performing a PCA on source and target independently, we restrict our analysis to a low-rank view of source and target data – which provides a first step filtering. Finally, although there are similarities in the maximization procedures from Equations Supp 11 and Supp 57, the product of our maximization procedure gives geometrical weights, and not directly the scores used in the regression. Although we maximize the same objective function, the constraints are different, which would make the final vectors surely different.
We believe our approach to be better suited for our specific problem for several reasons. First because it uses a low-rank representations of source and target. As shown in Figure 1 of main text, the kernel matrices $K_s$ and $K_t$ contain larger values than $K_{st}$ which would increase signal-to-noise ratio. Our sample-size is small – compared to single cell studies at least – and penalization is expected to focus on important signal. Second, our approach gives us a direct access to the geometric components (PV) which we can analyze to understand the making of the common signal. Finally, using PVs allow us to interpolate and get a projection on a single component that would be shared across source and target.

#### Supp 10 Algorithm workflow

**Algorithm 1** TRANSACT

**Require:** source data $\mathbf{X}_s$, target data $\mathbf{X}_t$, number of *domain-specific factors* $d_s$ and $d_t$, p.s.d. kernel $K$, number of *principal vector d.*

$\mathbf{K}_s \leftarrow$ source kernel matrix.
$\mathbf{K}_t \leftarrow$ target kernel matrix.
$\mathbf{K}_{st} \leftarrow$ source-target kernel matrix.
$\alpha^s \leftarrow$ Kernel Principal Components of source (from $K_s$).
$\alpha^t \leftarrow$ Kernel Principal Components of source (from $K_t$).
$\mathbf{M}^K \leftarrow \alpha^s C_{n_s} K_{st} C_{n_t} \alpha^t T.$
$\beta^s \Sigma \beta^t \leftarrow d$-rank SVD of $\mathbf{M}^K$, i.e. $\mathbf{M}^K \approx \beta^s \Sigma {\beta^t}^T$.
$\mathbf{F}^{st} \leftarrow \left[F^{st}(0), F^{st}(0.01), .., F^{st}(1)\right]$ defined as in Theorem Supp 6.6.
**for** $q \leftarrow 1$ to $d$ **do**
  $\mathbf{S}_q \leftarrow \left[\mathbf{F}^{st}[0]_{1:n_s,q}, \mathbf{F}^{st}[0.01]_{1:n_s,q}, .., \mathbf{F}^{st}[1]_{1:n_s,q}\right]^T$
  $\mathbf{T}_q \leftarrow \left[\mathbf{F}^{st}[0]_{n_s:n_s+n_t,q}, \mathbf{F}^{st}[0.01]_{n_s:n_s+n_t,q}, .., \mathbf{F}^{st}[1]_{n_s:n_s+n_t,q}\right]^T$
  $D_q \leftarrow \{D(S_q[0], T_q[0]), D(S_q[0.01], T_q[0.01]), .., D(S_q[1], T_q[1])\}.$
  $\tau_q^* \leftarrow \text{argmin}_\tau D_q.$
**end for**
$\mathbf{F} \leftarrow \left[\Phi_1(\tau_1), \Phi_2(\tau_2), .., \Phi_d(\tau_d)\right]$
$\tau^* \leftarrow \left[\tau_1^*, .., \tau_q^*\right].$
$X_s^{proj} \leftarrow \mathbf{F}^s t\left[\tau^*\right]_{1:n_s}.$
$X_t^{proj} \leftarrow \mathbf{F}^s t\left[\tau^*\right]_{n_s:n_s+n_t}.$
Train a regression model on $X_s^{proj}$
Apply it on the projected target data $X_t^{proj}$

### Supp 11 Glossary

| **Notation** | **Meaning** |
|---|---|
| $\mathbb{R}$ | Real numbers |
| $\mathbb{N}$ | Integers |
| $\|\| \cdot \|\|_F$ | Frobenius norm |
| $I_n$ | Identity matrix of size $n$. |
| $1_n$ | Vector of size $n$ with only ones. |
| $\mathcal{X}_s$ | Source data |
| $\mathcal{Y}_s$ | Source labels |
| $\mathcal{X}_t$ | Target data |
| $n_s, n_t$ | Number of source and target samples. |
| $p$ | Number of genes (features) |
| $d_s, d_t$ | Number of source and target principal components (NLPCs). |
| $d$ | Number of principal vectors pairs (and consensus features). |
| $\mathcal{F}(\mathbb{R}^p, \mathbb{R})$ | Spaces of functions from $\mathbb{R}^p$ to $\mathbb{R}$. |
| $K$ | Kernel functions ($\mathbb{R}^p \times \mathbb{R}^p \mapsto \mathbb{R}$). |
| $\gamma$ | Scaling factor of RBF kernel. |
| $\mathcal{H}$ | Feature space induced by $K$. |
| $\langle \cdot, \cdot \rangle_{\mathcal{H}}$ | Hilbert norm associated to K. |
| $\varphi$ | Feature map induced by $K$. |
| $K_s, K_t$ | Source and target kernel matrices |
| $K_{st} \in \mathbb{R}^{n_s \times n_t}$ | Kernel matrix between source and target samples. |
| $\mu_s$ | Mean source embedding. |
| $\widetilde{\varphi}_s$ | Feature map induced by $K$ translated by $\mu_s$. |
| $\mu_t$ | Mean target embedding. |
| $\widetilde{\varphi}_t$ | Feature map induced by $K$ translated by $\mu_t$. |
| $C_n$ | Centering matrix of size $n \in \mathbb{N}^*$. |
| $\alpha^s \in \mathbb{R}^{d_s \times n_s}$ | Sample importance scores of source NLPCs. |
| $\alpha^t \in \mathbb{R}^{d_t \times n_t}$ | Sample importance scores of target NLPCs. |
| $s_1, .., s_d$ | Source principal vectors (PV). |
| $t_1, .., t_d$ | Target principal vectors (PV). |
| span | Linear subspace generated by a family of vectors. |
| $\mathbf{M}^K \in \mathbb{R}^{d_s \times d_t}$ | Cosine similarity matrix between source and target NLPCs. |
| $\beta^s \in \mathbb{R}^{d_s \times d}$ | Left singular vectors of $\mathbf{M}^K$. |
| $\beta^t \in \mathbb{R}^{d_t \times d}$ | Right singular vectors of $\mathbf{M}^K$. |
| $\Sigma$ | Diagonal matrix with top $d$ singular values of $\mathbf{M}^K$. |
| $\rho^s \in \mathbb{R}^{d \times n_s}$ | Sample importance scores of source PVs. |
| $\rho^t \in \mathbb{R}^{d \times n_t}$ | Sample importance scores of target PVs. |
| $\theta_1, ..\theta_d$ | Principal angles between source and target PVs. |
| $\mathcal{S}^s, \mathcal{S}^t$ | Source data projected on source and target PVs. |
| $\mathcal{T}^s, \mathcal{T}^t$ | Target data projected on source and target PVs. |
| $\Gamma$ and $\xi$ | Angular interpolation functions. |
| $\phi_1, ..\phi_q$ | Interpolation between each pair of PVs. |
| $\tau = [\tau_1, .., \tau_d]$ | Interpolation times for each PV pair. |
| $\mathbf{F}^{st}(\tau)$ | Source and target data projected on interpolation at time $\tau$. |
| $\tau^* \in [0,1]^d$ | Optimal interpolation time for each PV pair. |
| $\mathcal{O}_1, .., \mathcal{O}_d$ | Offset contribution of each consensus feature. |
| $\mathcal{L}_{q,k}$ | Linear contribution of gene $k$ in consensus feature $q$. |
| $\mathcal{I}_{q,k,l}$ | Contribution of interaction between genes $k$ and $l$ to consensus feature $q$. |
